# Microbial Reduction of Metal-Organic Frameworks Enables Synergistic Chromium Removal

**DOI:** 10.1101/318782

**Authors:** Sarah K. Springthorpe, Christopher M. Dundas, Benjamin K. Keitz

## Abstract

Microbe-material redox interactions underpin many emerging technologies, including bioelectrochemical cells and bioremediation. However, commonly utilized material substrates, such as metal oxides, suffer from a lack of tunability and can be challenging to characterize. In contrast, metal-organic frameworks, a class of porous materials, exhibit well-defined structures, high crystallinity, large surface areas, and extensive chemical tunability. Here, we report that metal-organic frameworks can support the growth of the electroactive bacterium *Shewanella oneidensis.* Specifically, we demonstrate that Fe(III)-containing frameworks, MIL-100 and Fe-BTC, can be reduced by the bacterium via its extracellular electron transfer pathways and that reduction rate/extent is tied to framework structure, surface area, and particle morphology. In a practical application, we show that cultures containing *S. oneidensis* and reduced frameworks can remediate lethal concentrations of Cr(VI), and that pollutant removal exceeds the performance of either component in isolation or bioreduced iron oxides. Repeated cycles of Cr(VI) dosing had little effect on bacterial viability or Cr(VI) adsorption capacity, demonstrating that the framework confers protection to the bacteria and that no regenerative step is needed for continued bioremediation. In sum, our results show that metal-organic frameworks can serve as microbial respiratory substrates and suggest that they may offer a promising alternative to metal oxides in applications seeking to combine the advantages of bacterial metabolism and synthetic materials.

## MAIN TEXT

Redox-active minerals support the growth of metal-reducing and oxidizing microorganisms in anaerobic environments*(1)*. These microbe-mineral interactions play a key role in biogeochemical processes, including the nitrogen and carbon cycles*(2, 3)*, as well as nascent technologies like microbial fuel cells*(4)* and bioremediation*(5)*. For many of these applications, iron oxides are commonly employed as low-cost and widely available substrates for microbial reduction. However, iron oxides are significantly limited in structural diversity and only a fraction can be prepared under laboratory conditions*(6)*. Moreover, even relatively accessible structures, such as ferrihydrite (FeOOH·0.4H_2_O), are recalcitrant to structural characterization. Because applications like bioremediation depend on both microbial physiology and material characteristics, synthetic materials may offer an opportunity to more fully leverage microbial metabolism.

One emerging alternative to metal oxides are metal-organic frameworks. Relative to metal oxides, these materials generally exhibit higher surface areas, superior crystallinity, and can also possess oxide-like redox-activity*(7)*. More importantly, metal-organic frameworks facilitate the development of highly tunable structure-function relationships since properties such as pore size, linker identity, and metal node can be varied independently of one another*(8)*. As a result, they have received significant interest across several fields, including gas separations*(9)*, catalysis*(10)*, drug delivery*(11)*, and environmental remediation*(12)*. Despite their advantages over metal oxides and potential presence in nature*(13)*, metal-organic frameworks have not been examined as substrates for the growth of metal-reducing or oxidizing microorganisms. Due to their high surface areas and redox-active nature, we hypothesized that metal-organic frameworks could support microbial growth and potentially augment biological/material applications that rely upon biologically driven redox transformations.

Here, we demonstrate that iron-based metal-organic frameworks can serve as respiratory electron acceptors for the metal-reducing bacterium *Shewanella oneidensis* MR-1 (Figure 1). We selected Fe-BTC (BTC = 1,3,5-benzenetricarboxylate), Fe_3_O(BTC)_2_(OH)•nH_2_O (MIL-100), and Fe_3_O[(C_2_H_2_(CO_2_)_2_]_3_(OH)•nH_2_O (MIL-88A) for our growth and reduction assays, as their nodes are well-precedented MR-1 substrates (namely, Fe(III)), the frameworks are water-stable, and they span a range of morphologies. We observed that bacterial growth and metal reduction rate are intimately tied to framework structure, crystallinity, and surface area. Similar to growth on Fe(III) oxides, reduction was also governed by a known metal reduction pathway (MtrCAB) in *S. oneidensis* (Figure 1a) and influenced by the presence of soluble redox shuttles (flavins). Finally, in a potential application, we show that MR-1 and metal-organic frameworks synergistically facilitate bioremediation of Cr(VI). We found that microbial reduction of framework-bound Fe(III), especially in MIL-100, could generate high levels of redox-active Fe(II) that increased Cr(VI) reduction and adsorption to frameworks. Notably, Cr(VI) removal rates and capacities significantly exceeded those of biotic metal oxides, abiotic frameworks, and *S. oneidensis* alone. When operated together, the combination of MR-1 and MIL-100 also protected the bacteria from repeated challenges with lethal concentrations of Cr(VI) and required no separate regenerative steps. Overall, our results highlight the ability of metal-organic frameworks to support bacterial growth in a structure dependent manner and demonstrate how the advantages of these materials can be co-opted for heavy metal adsorption when coupled to bacterial metabolism.

**Figure 1.**
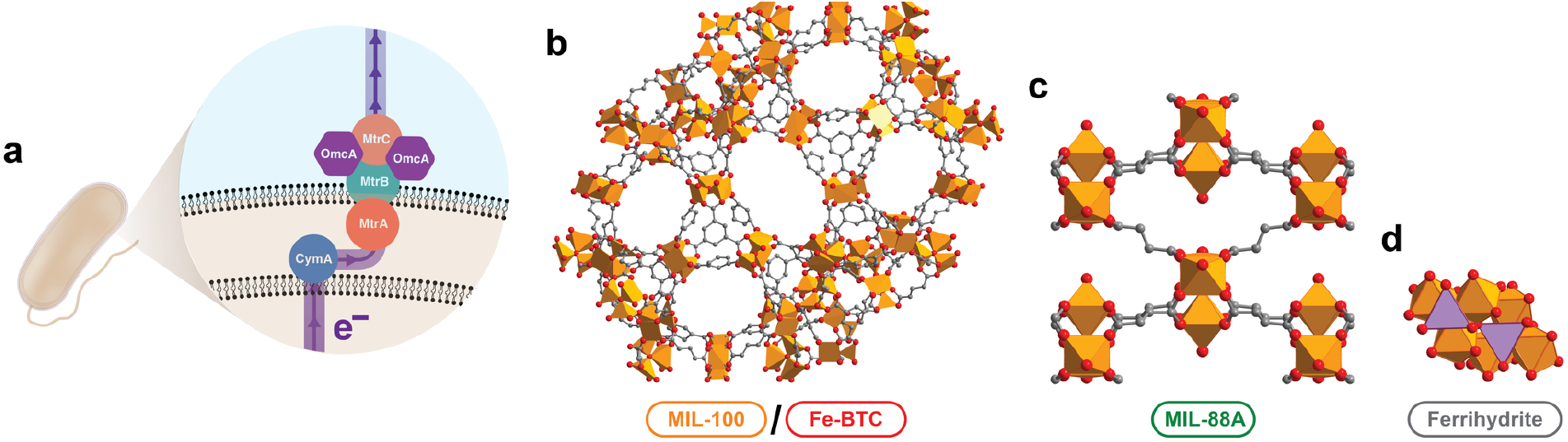
*S. oneidensis* MR-1 metal reduction pathway and metal-organic framework structure. (a) The MtrCAB pathway by which *S. oneidensis* reduces both soluble and insoluble metal species*(1)*. (b-d) The crystal structure of MIL-100/Fe-BTC (b, *(45, 46)),* MIL-88A (c, *(47)*), and ferrihydrite (d, *(24)*). Fe-BTC does not have a known crystal structure (due to its amorphous nature), but it is analogous to MIL-100.

## Results

### *S. oneidensis* MR-1 grows on metal-organic frameworks

First, we examined whether MR-1 could use iron-based metal-organic frameworks as respiratory substrates for cell growth. We prepared the metal-organic frameworks MIL-100, Fe-BTC, and MIL-88A and the commonly utilized *Shewanella* growth substrate ferrihydrite to draw comparisons with iron oxides. To monitor the growth of MR-1, each material was suspended in *Shewanella* Basal Medium (SBM) containing lactate as a carbon source and inoculated with anaerobically pregrown MR-1. Colony forming units (CFU) from dilutions of each material-bacteria suspension were used to measure cell counts. With MIL-100, Fe-BTC, and ferrihydrite suspensions, we observed comparable increases in cell concentration after 12 hours that stayed constant up to 24 hours (Figure 2a). In contrast, cell concentration in the MIL-88A suspension significantly decreased over a 24-hour period. As cell counts remained relatively constant in electron acceptor-free and *MIL-88A-Escherichia coli* controls (Figure S1), our results indicate that MIL-100 and Fe-BTC support MR-1 growth while MIL-88A causes Shewanella-specific cytotoxicity.

**Figure 2.**
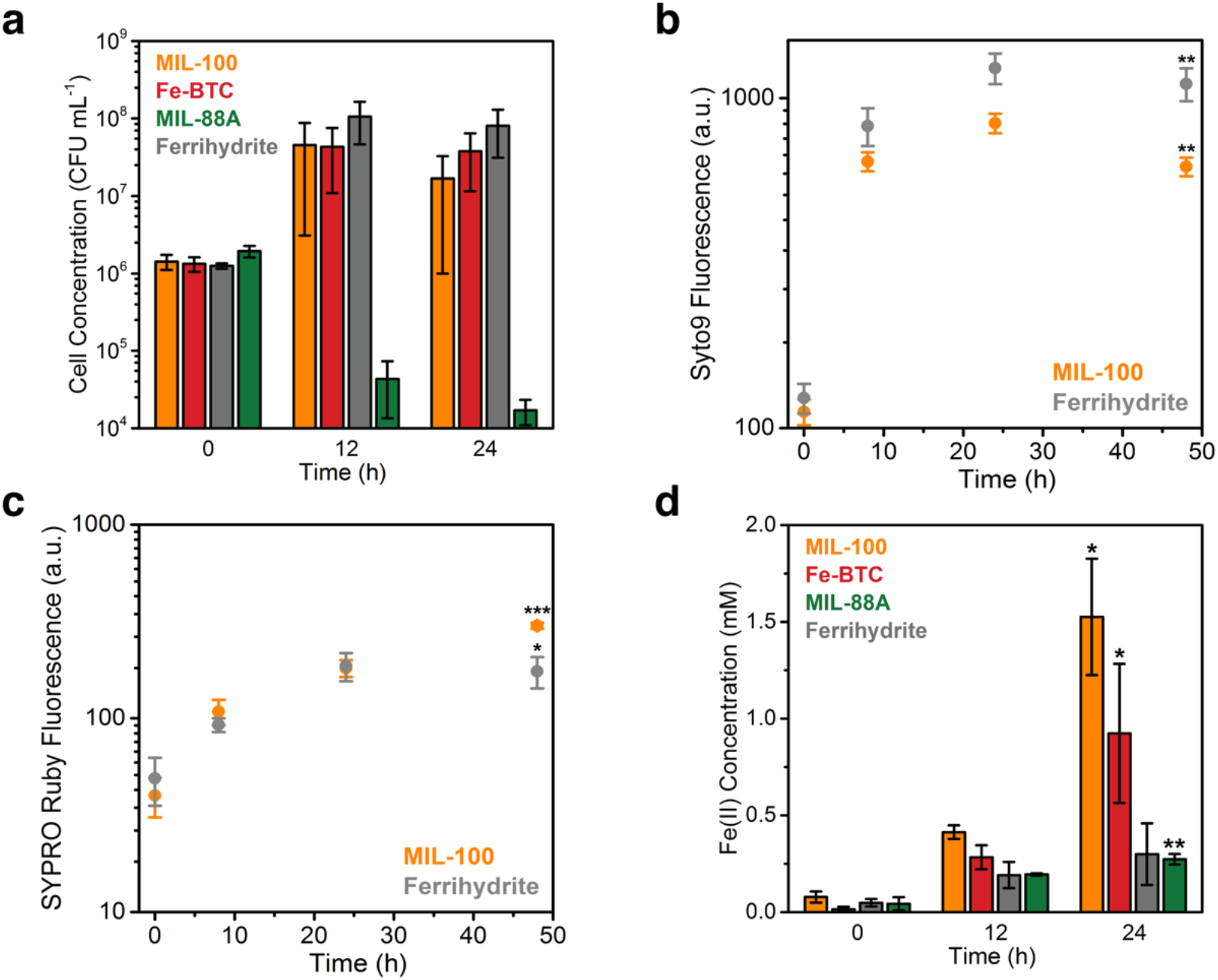
Growth of *S. oneidensis* MR-1 on metal-organic frameworks. (a) CFU counts and Fe(III) reduction of MR-1 (inoculating OD_600_=0.002) grown on MIL-100, Fe-BTC, MIL-88A, and ferrihydrite over 24 h. Data show mean ± S.E. for three independent biological replicates. (b) Measure of nucleic acids (Syto™ 9) for MR-1 (inoculating OD_600_=0.02) grown on MIL-100 and ferrihydrite over 48 h. Data show mean ± S.E. for four independent biological replicates. (c) Measure of biofilm matrix (SYPRO™ Ruby) for MR-1 (inoculating OD_600_=0.02) grown on MIL-100 and ferrihydrite over 48 h. Data show mean ± S.E. for four independent biological replicates. Figures (b) and (c) were collected in the same experiment. (d) Fe(III) reduction of MIL-100, Fe-BTC, MIL-88A, and ferrihydrite during growth of MR-1 over 24 h (inoculating OD_600_=0.002). Data show mean ± S.D. for three independent biological replicates. Figures (a) and (d) were collected in the same experiment. For Figures (b), (c), and (d), statistical comparisons were made between the first and last time points for each material using an unpaired two-tailed t-test; *P < 0.05, **P < 0.01, ***P < 0.001.

To further assess whether MR-1 was growing on the metal-organic frameworks, we used fluorescent dyes to measure other indications of biomass, such as increases in cellular DNA (Syto™ 9, ThermoFisher Scientific) and in biofilm matrix proteins (SYPRO™ Ruby, ThermoFisher Scientific). For both MIL-100 and ferrihydrite suspensions containing MR-1, Syto™ 9 fluorescence increased and rapidly plateaued to relatively similar extents after ~8 h (Figure 2b). Likewise, the SYPRO™ Ruby signal increased at a comparable rate for both material-bacteria suspensions (Figure 2c). We also used the auto-fluorescence of secreted flavins (flavin mononucleotide, riboflavin, flavin adenine dinucleotide) to quantify differences in biomass supported by the frameworks*(14)*. Auto-fluorescence increased concomitantly with CFU counts from MIL-100 suspensions and with optical density in planktonic cultures growing on fumarate (Figure S2). With the exception of MIL-88A, we observed a similar increase in auto-fluorescence when MR-1 was grown on each framework and ferrihydrite. Together, these results corroborate our CFU counts and demonstrate that MR-1 is able to grow on some iron-based metal-organic frameworks. Moreover, in the absence of material toxicity, biomass accumulation appears relatively invariant to the framework used and comparable to growth with a model iron oxide (ferrihydrite).

Over the course of growth quantification by CFU counts, Fe(II) levels were simultaneously monitored for each material-bacteria suspension. Across all experiments, Fe(II) levels did not change in abiotic controls, but did increase in conjunction with cell growth. Increases in Fe(II) concentrations were significantly different based on material identity (Figure 2d), despite similarities in cell counts between MIL-100, Fe-BTC, and ferrihydrite suspensions. After 24 hours, MIL-100 suspensions exhibited the highest increase in Fe(II) concentration (1.5 ± 0.3 mM) and Fe-BTC the second-highest (0.92 ± 0.4 mM). Fe(II) levels only slightly increased over 24 hours for MIL-88A (0.30 ± 0.03 mM) and ferrihydrite (0.30 ± 0. 2 mM). Similar results were observed for Fe(II) levels measured simultaneously with Syto™ 9 and SYPRO™ Ruby measurements (Figure S3). Although final Fe(II) concentrations were different for each material tested, our results are in agreement with previous *Shewanella* growth studies on soluble/insoluble iron substrates, which showed that cell yields were similar regardless of electron acceptor used(15).

### Microbial reduction rate is dependent on framework structure

As biological reduction of each metal-organic framework differed substantially, we next analyzed how material structure affected the rate of Fe(III) reduction by MR-1. Significant amounts of Fe(II) were observed when MR-1 was grown on MIL-100 and Fe-BTC (Figure S4). For Fe-BTC, MR-1 reduced 1. 9±0.3 mM Fe(III) before plateauing after 72 h. The largest extent of reduction was observed with MIL-100, with Fe(II) concentrations reaching 5.1 ±0.6 mM Fe(II) after 144 h. Assuming all Fe(III) in MIL-100 was reduced, the theoretical maximum Fe(II) concentration would be 15 mM. Thus, our results indicate that MR-1 can access approximately a third of the Fe(III) in the MIL-100 structure. As expected from our growth experiments, a minimal amount of reduction was observed when a suspension of MIL-88A was inoculated with MR-1. The above results correspond to rates of 47.4±0.2 μM h^-1^ for MIL-100, 22.3±3.4 μM h^-1^ for Fe-BTC, and 3.8±1.3 μM h^-1^ for MIL-88A over the linear range of Fe(III) reduction (Figure 3a). For comparison, MR-1 grown on ferrihydrite showed a reduction rate of 5.2±1.7 μM h^-1^, which is consistent with previous reports when normalized by inoculating OD_600_(16). These data confirm that Fe(III) reduction rate strongly depends on framework structure.

**Figure 3.**
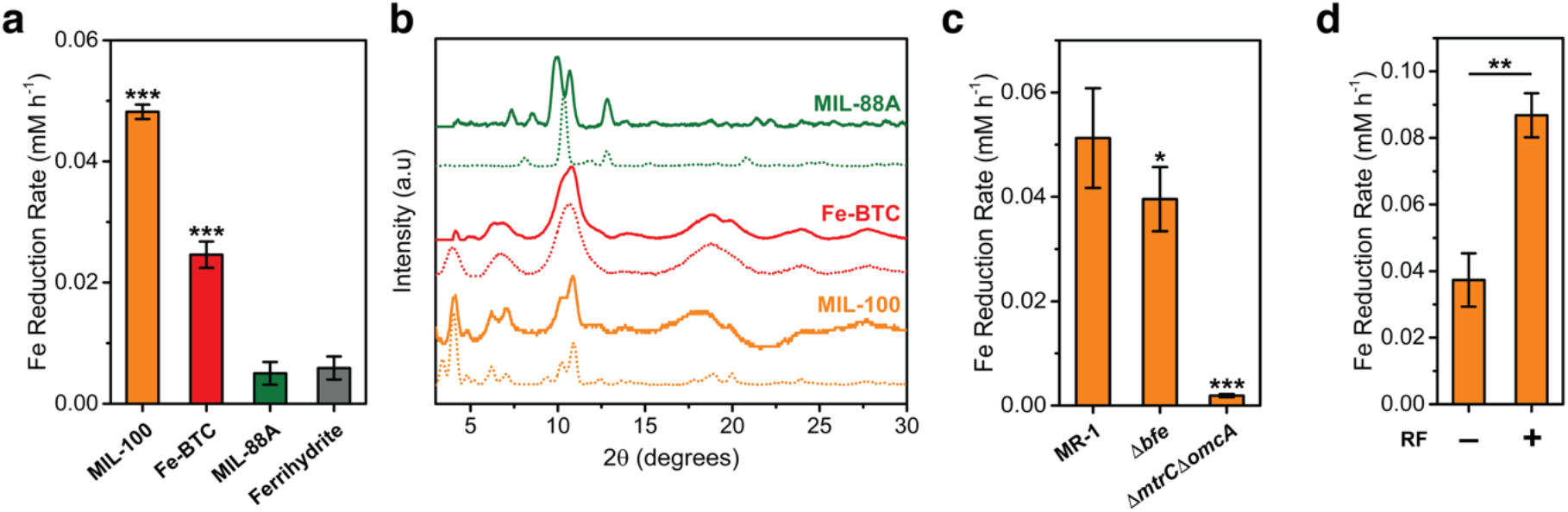
*S. oneidensis* reduction rates of metal-organic frameworks. (a) Reduction rates of MIL-100, Fe-BTC, MIL-88A, and ferrihydrite over 96 h. Data shown are mean ± S.D. for three independent biological replicates. Reduction rates that statistically differ from ferrihydrite are indicated; ***P< 0.001, ANOVA with post-hoc Dunnett’s test. (b) As-synthesized (dashed lines) and post-reduction (solid lines) PXRD patterns of MIL-100, Fe-BTC, and MIL-88A. (c) MIL-100 reduction rates of MR-1, *Δbfe,* and *>ΔmtrC>ΔomcA* grown over 48 h. Data shown are mean ± S.D. for six independent biological replicates. Reduction rates that statistically differ from MR-1 are indicated; *P < 0.05, ***P < 0.001, ANOVA with post-hoc Dunnett’s test. (d) Reduction rates of MIL-100 by MR-1 with and without exogenous 10 μM riboflavin supplementation. Data show mean ± S.D. for three independent biological replicates. **P < 0.01, unpaired two-tailed t-test.

Next, we investigated how the structural and morphological differences between the frameworks contributed to the observed variation in iron reduction rates. MIL-100 and Fe-BTC are closely related in molecular structure, with MIL-100 being more crystalline. Interestingly, iron oxides with higher degrees of crystallinity are typically associated with slower Fe(III) reduction rates across the *Shewanella* genus*(17, 18)*. However, the extent of Fe(III) reduction by metal-reducing bacteria is also dependent on the interplay between particle size, extent of aggregation, and surface area*(19, 20)*. Furthermore, it has been found that the interfacial contact between outer membrane cytochromes and iron oxides impacts reduction rates, with smaller particles unable to provide adequate contact for reduction*(21)*. To determine if similar material features influenced the biological reduction of metal-organic frameworks, we examined the morphology of MIL-100, Fe-BTC, and MIL-88A using scanning electron microscopy (SEM). SEM images showed that MIL-100 and Fe-BTC were highly aggregated, while MIL-88A was comprised of small, well-defined crystals approximately the same size as MR-1 (Figure S5). Based on its morphology, MIL-88A is unlikely to support biofilm development*(22, 23)*. Although they are morphologically similar, MIL-100 and Fe-BTC vary significantly in accessible surface area. MIL-100 had the largest Langmuir surface area (2188 m^2^g^-1^) followed by Fe-BTC (1512 m^2-1^), ferrihydrite (311 m^2-1^, *(24))* and MIL-88A (130 m^2-1^) (Table S1). These surface areas closely track our measured Fe(III) reduction rates. Overall, our results are consistent with previous studies of MR-1 growth on iron oxides and show that biotic metal-organic framework reduction is governed through a combination of particle morphology, framework structure, and accessible surface area.

### Stability of metal-organic frameworks in the presence of MR-1

Next, we asked if Fe(III) reduction and metabolic activity from MR-1 negatively impacted the solution stability of the frameworks. Framework stability under typical culture conditions in the absence of bacteria was confirmed by monitoring Fe(III) and fumarate leaching, as well as PXRD (Figure S6). MIL-100 and Fe-BTC maintained structural integrity while MIL-88A exhibited a slight, but detectable, change in structure. From PXRD patterns collected post-reduction, we found that MIL-88A eventually decomposed under biotic conditions after 48 h (Figure 3b). In contrast, characteristic MIL-100 peaks were discernable after exposure to MR-1. This result indicates that MIL-100 remains structurally intact and crystalline over the course of our experiments. Additionally, we observed that MIL-100 and Fe-BTC changed color following reduction by MR-1, suggesting Fe(II) was contained in the framework (Figure S7). This was further evidenced by measured Fe(II) in the washed samples of biotic MIL-100 and Fe-BTC. Nonetheless, the bacteria were able to solubilize some iron in these frameworks. Similar to observations with iron oxides that are partially solubilized upon microbial reduction, our results indicate that MR-1 can both directly reduce Fe(III) in MIL-100 and partially remove Fe(II/III) from the framework structure while maintaining framework stability(25).

### The MtrCAB pathway controls metal-organic framework reduction

The MtrCAB pathway in MR-1 is largely responsible for extracellular electron transfer and interfacing with insoluble metal oxides*(16)*. Specifically, the outer-membrane cytochromes MtrC and OmcA enable the terminal electron transfer step onto both soluble and insoluble metal species*(26)*. Without these proteins, *S. oneidensis* shows attenuated respiration onto iron and graphene oxides*(27)*. Consistent with these results, an outer-membrane cytochrome deficient strain, *DmtrCDomcA,* showed minimal Fe(III) reduction relative to MR-1 (Figure 3c, Figure S8). *E. coli* MG1655, which does not possess a homolog to MtrC or OmcA and lacks electroactivity, was also unable to reduce MIL-100 or Fe-BTC (Figure S9). Together, these results confirm that the same biological pathways responsible for iron oxide reduction by *S. oneidensis* also play a key role in its reduction of metal-organic frameworks.

### Exogenous flavins improve Fe(III) reduction in metal-organic frameworks

In addition to the MtrCAB pathway, MR-1 uses soluble redox shuttles (flavins) to transfer reducing equivalents to metal oxides, with exogenously supplied flavins accelerating the rate of Fe(III) reduction*(28)*. Similarly, we found that addition of 10 μM riboflavin to MR-1 cultures containing MIL-100 increased Fe(III) reduction rates and the total amount of Fe(II) (Figure 3d, Figure S10). In the presence of excess flavins, MR-1 could access approximately half of the Fe(III) in the MIL-100 framework. When grown on MIL-100, a strain lacking flavin export machinery *(Dbfe)* showed a slight, but significant, decrease in Fe(III) reduction rate relative to MR-1 (Figure 3c). These results are consistent with previous studies where the *Dbfe* strain exhibited a similar decrease in Fe(III) reduction compared to MR-1 when both were grown on ferrihydrite*(29)*. Overall, our data indicate that flavins are important contributors to Fe(III) reduction in metal-organic frameworks.

### Cr(VI) adsorption by microbially-reduced metal-organic frameworks

Having determined that microbial metabolism can interface with metal-organic frameworks, we tested whether this pairing could be applied towards redox transformations. These transformations, especially the reduction of iron oxides and release of reactive Fe(II), play a major role in the aqueous transport of heavy metal contaminants and can be performed by a variety of metal-reducing bacteria, including MR-1*(30)*. Moreover, electroactive bacteria can directly reduce a variety of heavy metal species*(30)*, increasing their utility for redox based applications. As a result, metal-reducing bacteria, acting in conjunction with iron oxides, have shown promise for the bioremediation of redox-controlled toxic contaminants, such as CrO_4_^-2^ or Cr_2_O_7_^-2^*(31)*. Alternatively, standard inorganic adsorbents, including activated carbons and metal-organic frameworks can be used for the remediation of these contaminants. For example, some metal-organic frameworks have been used for Cr(VI) adsorption, but generally show slow adsorption kinetics and limited capacities*(32–34)*. A more recent study found that Cr(VI) adsorption to a metal-organic framework could be dramatically improved by adding exogenous Fe(II) *(35)*. This result implies that, similar to the case with iron oxides, the presence of Fe(II) can enhance reductive adsorption of Cr(VI). Thus, we hypothesized that microbial reduction of Fe(III)-containing frameworks could exhibit a synergistic effect on Cr(VI) removal (Figure 4a).

**Figure 4.**
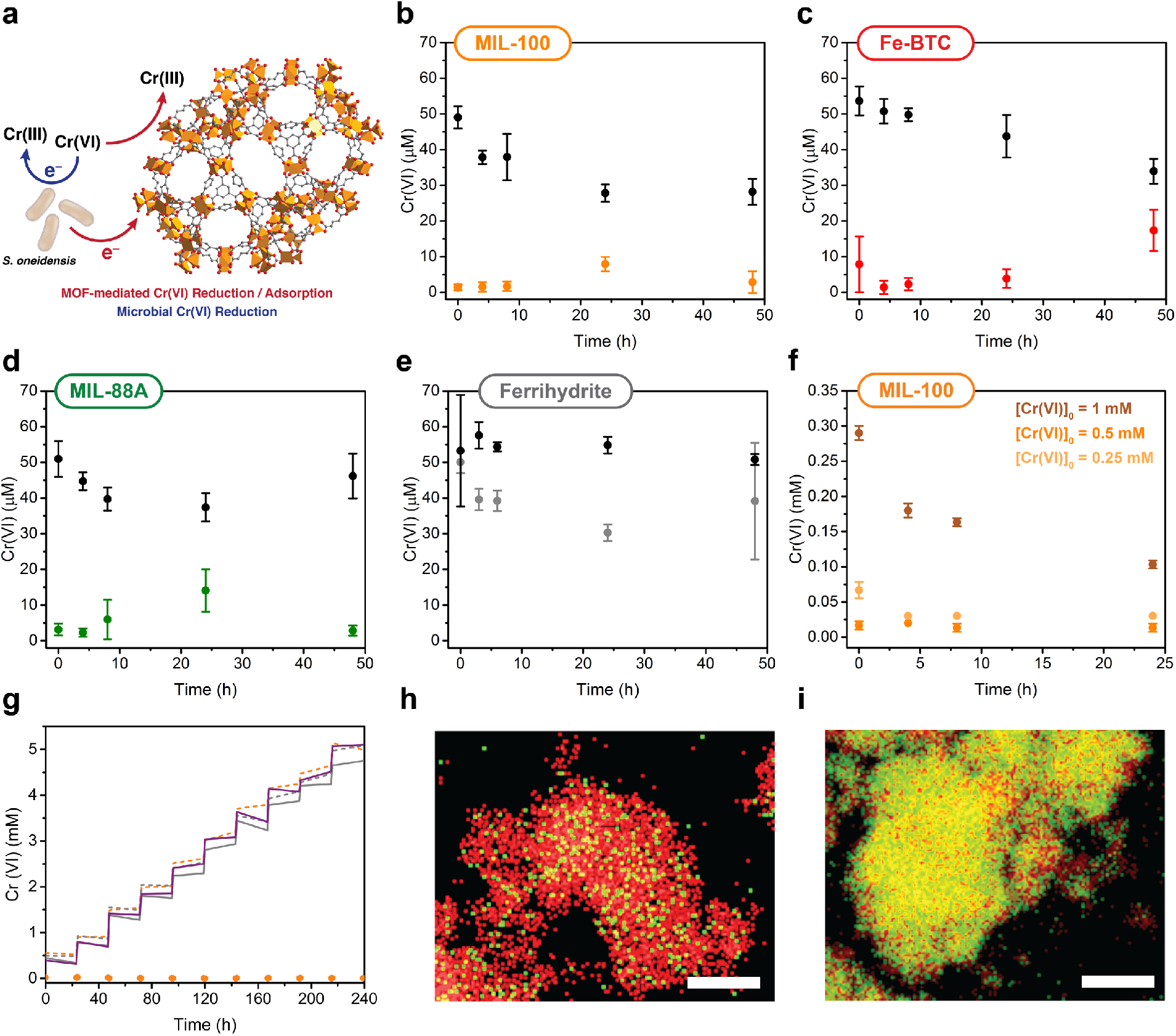
Cr(VI) removal in MR-1 reduced metal-organic frameworks. (a) Generalized scheme for Cr(VI) reduction by both MR-1 and a reduced metal-organic framework. (b-e) Cr(VI) removal by MR-1-reduced (b) MIL-100, (c) Fe-BTC, (d) MIL-88A, and (e) ferrihydrite when challenged with 70 μM Cr(VI). Abiotic samples for each material are shown in black. (f) Cr(VI) removal by MR-1-reduced MIL-100 when challenged with 0.25 mM, 0.5 mM, and 1 mM Cr(VI). (g) Cr(VI) removal in MR-1-reduced MIL-100 (orange), ferrihydrite (grey) and fumarate (purple) when challenged with ten cycles of 0.5 mM Cr(VI). Abiotic samples are shown as dashed lines. (h-i) EDX element mapping of (h) abiotic cycled MIL-100 and (i) MR-1-reduced and cycled MIL-100. Fe is shown in red and Cr is shown in green. Scale bars are 1 μm. Data show mean ± S.D. for three independent biological replicates.

In initial experiments, we grew MR-1 on each metal-organic framework for 24 hours and then challenged the culture with a single dose of Cr(VI). At initial Cr(VI) concentrations of 70 μM, all frameworks showed immediate removal of soluble Cr(VI) below the detection limit of the diphenylcarbazide (DPC) assay (Figure 4b-d). Abiotic controls and biotic ferrihydrite (Figure 4e) showed minimal and slow Cr(VI) adsorption. In the case of MIL-100 and Fe-BTC, we found that Fe(III) reduction continued to take place after Cr(VI) addition, indicating that MR-1 was still viable under these conditions. Even though MIL-88A showed minimal Fe(III) reduction in the presence of MR-1, it was still enough to reductively adsorb the relatively small initial dose of Cr(VI). These results are a preliminary indication that there is a synergistic interaction between the frameworks and MR-1 that accelerates Cr(VI) reduction and adsorption.

Next, we challenged MR-1 growing on MIL-100 with increasing concentrations of Cr(VI) up to 1 mM (Figure 4f). After dosing with 0.5 mM Cr(VI), MR-1 and MIL-100 reduced all detectable Cr(VI) within minutes. Under these conditions, the total solution concentration of Cr, as measured by inductively coupled plasma – mass spectrometry (ICP-MS), fell below 10 ppm within minutes and reached ~1 ppm after 24 hours (Figure S11). Furthermore, Fe(III) reduction continued to take place after Cr(VI) addition, suggesting that MR-1 can survive normally toxic Cr(VI) concentrations in the presence of MIL-100. At 1 mM Cr(VI), MR-1 on MIL-100 showed almost complete reduction, but over a much longer time period, indicating that this concentration is approaching the limit of toxicity for our system.

Finally, we sought to distinguish biological reduction of Cr(VI) from framework-mediated reductive adsorption. As mentioned above, MR-1 can directly reduce Cr(VI) through the MtrCAB pathway. Alternatively, MR-1 could mediate Cr(VI) reduction through MIL-100. Overall, we found that biotic samples in the absence of MIL-100 showed significantly slower Cr(VI) reduction kinetics and were more sensitive to the initial Cr(VI) dose (Figure S12). For example, MR-1 challenged with 500 μM Cr(VI) was only reduced ca. 100 μM after 24 hours and entered death phase, as measured by OD_600_ (Figure S13). These results are consistent with previous reports indicating that 200 μM Cr(VI) is the upper limit for MR-1 survivability*(36)*. We also measured reduced Cr(VI) adsorption kinetics for MR-1 grown on ferrihydrite. Notably, even at low initial Cr(VI) concentrations, biotic and abiotic ferrihydrite samples exhibited slower adsorption kinetics and capacities relative to all frameworks tested (Figure 4). Additionally, after MR-1 had reduced appreciable amounts of Fe(III) in ferrihydrite, there was not a corresponding decrease in Cr(VI), as would have been expected if Fe(II) alone was responsible for Cr(VI) reduction. Thus, our results support a synergistic interaction between MR-1 and MIL-100 that enhances Cr(VI) reduction/adsorption kinetics and capacity relative to abiotic frameworks and MR-1 growing on iron oxides.

### Repeated cycles of Cr(VI) with MR-1 reduced MIL-100

The majority of materials for Cr(VI) adsorption require a regeneration step using pH adjustments after saturation*(37)*. Based on our initial Cr(VI) adsorption experiments involving MR-1 and MIL-100, we predicted that viable cells could metabolically regenerate the framework for continued Cr(VI) reduction and adsorption. To determine and compare the recyclability of MR-1-reduced materials, samples containing either MIL-100, ferrihydrite, or fumarate were reduced by MR-1 for 24 h before being challenged with 10 cycles of [Cr(VI)]0=0.50 mM, once every 24 hours. For all additions to MR-1 growing on MIL-100, soluble Cr(VI) was removed below the limit of detection within minutes (Figure 4g). The total amount of Cr(VI) reduced by MR-1 and MIL-100 was 78.7±0.1 mg g^-1^ (Table S2), which represents a 125-fold increase over MIL-100 alone and compares favorably to other Cr(VI) adsorbents (Table S3). Moreover, when the maximum Cr(VI) capacity was tested, we found that 320.3±0.6 mg g^-1^ was reduced, accounting for 75% of the theoretical maximum, assuming all electrons from MR-1 lactate consumption are used for Cr(VI) reduction (Figure S14).

In contrast, biotic and abiotic treatments with fumarate and ferrihydrite showed no appreciable Cr(VI) adsorption and the amount of Cr(VI) in solution increased after each new dose. We also investigated whether a microbially-reduced soluble iron source could aid in removal of Cr(VI). Indeed, we found that MR-1 grown on Fe(III)-citrate enabled instantaneous reduction after repeated cycles of 0.50 mM Cr(VI) (Figure S15). However, no precipitate was formed upon Cr(VI) reduction. Similar results were found for abiotic reduction of Cr(VI) by FeCl2·4H_2_O, with no precipitate forming despite complete reduction. These results suggest that, under our pH operating conditions, Cr(III) can only be removed from solution in the presence of a sorbent material, such as MIL-100. After Cr(VI) cycling, we characterized MIL-100 using scanning transmission electron microscopy (STEM) and transmission electron microscopy (TEM). Abiotic and biotic samples were morphologically similar (Figure S16). In contrast, elemental mapping revealed that cycled abiotic MIL-100 was primarily composed of Fe with small amounts of Cr while cycled biotic MIL-100 contained significantly more Cr (Figure 4h-i). This result supports our solution-based measurements and confirms that metabolic reduction of MIL-100 significantly increases its overall capacity for Cr(VI) adsorption. Our cycling data suggests that MR-1 growing on fumarate or ferrihydrite is quickly overwhelmed after repeated challenges with Cr(VI). Indeed, we observed no increase in OD_600_ in the fumarate samples following the first addition of Cr(VI), implying that the cells had entered death phase (Figure S13). In contrast, we measured an increase in Fe(II) over the course of the cycling experiment when MR-1 was grown on MIL-100. As expected, Fe(II) concentrations decreased immediately after the addition of Cr(VI) but rebounded as viable cells continued to reduce Fe(III) (Figure S17). We did not measure a similar trend in Fe(III) reduction for MR-1 grown on ferrihydrite that was repeatedly challenged with Cr(VI) (Figure S17). We also verified the observed trends in Fe(III) reduction during Cr(VI) cycling by directly assaying cell viability. Following the tenth Cr(VI) addition, the MIL-100 sample formed a lawn after an aliquot was plated on LB agar and grown aerobically overnight, indicating viable bacteria remained after Cr(VI) cycling (Figure S18). In contrast, abiotic samples, biotic ferrihydrite samples, and biotic fumarate samples showed no growth on LB agar plates. Together, these results demonstrate that MIL-100 both effectively protects MR-1 from repeated challenges of cytotoxic Cr(VI) concentrations and synergistically enables Cr(VI) adsorption with rates and capacities that greatly exceed those of biotic metal oxide or abiotic metal-organic frameworks.

## Discussion

While previously limited to metal oxides, noble metals, and other electrode materials, we demonstrated that microbe-material redox interactions can be extended to metal-organic frameworks. The primary advantage of these materials is their high degree of synthetic tunability. By changing the metal node and/or linker identity, material structure and functionality can be radically altered. Although we examined only a small selection of frameworks, our results show that pairing these materials with electroactive microbes can actuate vastly different biological (e.g., cell growth, Fe(III) reduction rates) and chemical (e.g., Cr(VI) removal) responses.

We observed that iron-containing metal-organic frameworks can serve as growth substrates for *S. oneidensis.* Biomass measurements and quantitation of Fe(II) levels determined that these materials behave as respiratory electron sinks similarly to a model iron oxide (namely, ferrihydrite). Our genetic knockout and flavin-supplementation experiments further evidence that canonical extracellular electron tranfer pathways of *S. oneidensis* participate in Fe(III) reduction. Notably, reduction rates were greatly accelerated over ferrihydrite with the frameworks MIL-100 and Fe-BTC. Thus, these materials may find use as alternative insoluble substrates when studying electroactive physiology. To this end, metal-organic frameworks can be constructed from a variety of other biologically-relevant metal nodes, including manganese, chromium, and cobalt*(8)*. Conceivably, the organic linker or other framework components could also be modified to act as secondary electron acceptors. For example, frameworks that contain DMSO (Zn2-(4,4’-Ethyne-1,2-diyldibenzoate)2(DMSO)2]1.6H_2_O) *(35)*, porphyrins (PCN-222) *(39)*, or conductive linkers (Co2TTFTB; TTFTB = tetrathiafulvalene tetrabenzoate) *(40)* could potentially support microbial growth. The PCN-222 framework is of particular interest as the metal center of the porphyrin linker can be exchanged without altering the overall structure and it is highly stable in water*(39)*. Finally, pore size can also be tuned, potentially controlling the diffusion of flavins and other redox shuttles in the framework.

Despite their potential advantages, a major limitation of applying metal-organic frameworks as microbial growth substrates is their general instability under aqueous conditions. To counteract this, we examined water stable frameworks and also note that there are significant efforts to develop additional frameworks with enhanced water stability*(41)*. Nevertheless, the aqueous stability of a specific framework must be carefully scrutinized before it is selected as a substrate for microbial growth. For example, we found that stepanovite, a naturally-occurring metal-organic framework mineral*(13)*, could also support the growth of MR-1, but that this effect was due to dissolution of the framework (Figure S19). Potentially, utilization of water-stable frameworks possessing well-defined structures could reduce the impact of abiotic material effects, such as the phase or polymorph changes that frequently occur in metal oxides*(6)*. The effect of selected frameworks on bacterial viability should also be carefully assessed to ensure cellular functioning when desired. For example, MIL-88A is comprised of two known *S. oneidensis* substrates (Fe(III) and fumarate), but we unexpectedly observed a cytotoxic effect on *S. oneidensis* and not *E. coli.* Nonetheless, the synthetic tunability of metal-organic frameworks should position them as useful tools for probing material effects on microbial physiology and improving the design of applications that exploit these relationships.

In one potential application, we showed that MR-1 operates synergistically with MIL-100 to reduce and adsorb significant amounts of Cr(VI). Notably, the high adsorption capacity of MIL-100 shielded the bacteria from repeated Cr(VI) challenges while analogous materials (ferrihydrite) provided no protection. Additionally, since Fe(III) is continually reduced by MR-1, then reoxidized after Cr(VI) addition, the measured capacity of reduced MIL-100, corresponding to 320 mg g^-1^, is only limited by the amount of lactate in culture medium and represents a 523-fold increase relative to abiotic MIL-100. Our results with MIL-100 also suggest that other redox-active frameworks could see a similar increase in Cr(VI) adsorption capacity when combined with MR-1. Alternatively, frameworks could support the growth of metal-reducing bacteria to synergistically remediate other environmental pollutants, like U(VI) *(42)*, or could support syntrophic cocultures of different bacterial species. For example, iron oxides can serve as mediators of electron transfer between methanogens and *Geobacter sulfurreducens(43, 44)*; iron-based metal-organic frameworks, including MIL-100, could play a similar role. Finally, metal-organic frameworks may be exciting materials to include in microbial fuel cells since their large surface areas, porosity, and ability to support bacterial growth may translate into performance improvements in these bioelectrochemical devices*(4)*. Overall, metal-organic frameworks offer promise for optimizing a wide range of technologies underpinned by microbe-material redox interactions.

## Materials and Methods

### Synthesis of metal-organic frameworks and iron oxides

MIL-100 was synthesized by first dissolving H_2_BTC (1.68 g, 7.6 mmol) and NaOH (0.91 g, 22.8 mmol) in 24 mL H_2_O. This solution was added dropwise to FeC_2_4H_2_O (2.26 g, 11.4 mmol) dissolved in 97 mL H_2_O and stirred for 24 h at room temperature/ambient conditions. The precipitate was isolated, washed with water (3x at 25 °C for 12 h) and ethanol (1x at 25 °C for 12 h), and then dried at room temperature/ambient conditions*(45)*. Fe-BTC was synthesized by dissolving H3BTC (0.263 g, 1.2 mmol) and NaOH (0.15 g, 3.8 mmol) in 10 mL of H_2_O. This solution was added dropwise to FeCl3·6H_2_O (0.513 g, 1.9 mmol) dissolved in 10 mL H_2_O and stirred for 10 min at room temperature/ambient conditions. The precipitate was isolated, washed with water (3x at 25 °C for 12 h) and ethanol (1x at 25 °C for 12 h), and then dried at room temperature/ambient conditions*(46)*. MIL-88A was synthesized by stirring a solution of fumaric acid (0.97 g, 8.4 mmol) and FeCl_3_·6H_2_O (2.27 g, 8.4 mmol) in 42 mL of H_2_O for 1 h before transferring to a Teflon-lined steel autoclave (Parr). The reactor was heated at 65 °C for 12 h and then cooled down to room temperature. Precipitate from inside the reactor was subsequently washed with ethanol (3x at 25 °C for 12 h) and water (3x at 25 °C for 12 h), then dried at 120 °C for 10 h*(47)*. Ferrihydrite was synthesized by adding 1M NaOH dropwise to FeCl3·6H_2_O (5.4 g, 20.0 mmol) dissolved in 100 mL H_2_O until the pH reached ~7.5. The solid was isolated by centrifugation, washed with water (3x at 25 °C for 10 min), and lyophilized immediately following washing for 48 h(24). Stepanovite was synthesized by stirring red hematite (1.5 g, 9.5 mmol), NaOH (0.38 g, 9.5 mmol), and magnesium oxide (0.38, 9.5 mmol) in 30 mL of 10% w/v oxalic acid solution for 15 h. Single crystals were obtained by filtering and storing the solution at 4 °C for 24 h. Stepanovite powder used for MR-1 reduction was obtained by evaporating the solution after filtration using a rotary evaporator (85 °C, 45 rpm, low vacuum, *(13)*).

### Strains and culture conditions

All anaerobic cultures and experiments were performed inside a humidified Coy anaerobic chamber (3% H_2_/balance N2 atmosphere). *S. oneidensis* MR-1 was obtained from ATCC^®^ (700550™). Mutant strains, *DmtrCDomcA* and *Dbfe,* were generously provided by JA Gralnick (University of Minnesota, Minneapolis, MN). For anaerobic pregrowths, all strains were cultured in SBM buffered with 100 mM HEPES and supplemented with 0.05% casamino acids and 0.5% Wolfe’s mineral mix*(48)*. The medium was adjusted to a pH of ~7.2. For pregrowth, sodium lactate (20 mM) and sodium fumarate (40 mM) were added as the carbon source and an electron acceptor, respectively. Once the bacteria reached stationary phase (ca. 18 h), they were pelleted by centrifugation (6000 x g; 20 min) and washed with SBM two times. A final addition of SBM was added to dilute the cells to OD_600_=0.2.

### Growth of MR-1 on metal-organic frameworks and iron oxides

Growth of MR-1 on the metal-organic frameworks was measured by CFU counting, Syto™ 9 fluorescent dye, SYPRO™ Ruby fluorescent dye, and auto-fluorescence. Cultures containing 20 mM sodium lactate and either MIL-100, Fe-BTC, MIL-88A, or ferrihydrite ([Fe(III)]_0_=15 mM) in fresh SBM were prepared anaerobically and then inoculated with the washed MR-1. CFU/Fe(III) reduction studies used an inoculating OD_600_=0.002. Syto™ 9/SYPRO™ Ruby/Fe(III) reduction studies used an inoculating OD_600_=0.02 to increase the fluorescent signal. Abiotic samples were prepared with the materials in SBM with lactate but with no cells. The cultures were incubated anaerobically at 30 °C for up to 48 h. For CFU counting, an aliquot of the culture suspension was removed and serially diluted (10^0^-10^-7^). Each dilution (100 μL) was plated onto LB agar plates using sterile glass beads and incubated aerobically at 30 °C overnight. Individual colonies on each plate were counted and the initial concentration of MR-1 was calculated. For the staining of bacterial nucleic acids, 200 μL of the culture suspension was removed and pelleted before removing the supernatant. The pellet was resuspended in 100 μL of 0.85% NaCl solution, mixed with 2X Syto™ 9 solution in a 96 well plate, and incubated at room temperature for 15 minutes in the dark. Fluorescence was measured with an excitation/emission of 485/530 nm using a BMG LabTech CLARIOstar Monochromator Microplate Reader. For the staining of biofilm matrix, 100 μL of the culture suspension was removed and pelleted before removing the supernatant. The pellet was mixed with 200 μL of 1X SYPRO™ Ruby fluorescent dye in a 96 well plate. The plate was incubated in the dark at room temperature for 15 minutes before measuring fluorescence at an excitation/emission of 460/640 nm using a platereader. For the auto-fluorescence assay, 250 μL of the culture suspension was removed, gently spun down to pellet the material, and the supernatant was transferred to a 96-well plate. Fluorescence was measured with an excitation/emission of 485/528 nm using a platereader.

### Reduction of metal-organic frameworks and ferrihydrite

Reduction of the metal-organic frameworks and ferrihydrite by MR-1 was tested by measuring Fe(II) concentrations in framework-bacteria suspensions over time. Suspensions containing 20 mM sodium lactate and either MIL-100, Fe-BTC, MIL-88A, or ferrihydrite ([Fe(III)]0=15 mM) in fresh SBM were prepared anaerobically and then inoculated with the washed MR-1 to OD_600_=0.002. Abiotic samples were prepared with the materials in SBM with lactate but with no cells. The suspensions were incubated anaerobically at 30 °C for 144 h. At each time point, the material substrates were gently shaken to fully suspend all materials before removing an aliquot from the culture. Each aliquot was mixed with 6 M HCl in a 1:1 ratio, until full dissolution of the framework or oxide occurred. Then, the sample was diluted with 1 M HCl in a 1:1 ratio and analyzed for Fe(II) using the ferrozine assay*(49)*. The ferrozine assay was performed by mixing 15 μL of the diluted sampled with 235 μL of ferrozine solution in a 96-well plate. Absorbance was measured at 562 nm using a microplate reader. Ferrozine solution consisted of ferrozine (45 mg, 87 μmol) and ammonium acetate (22.5 g, 0.29 mol) in 45 mL H_2_O. A similar protocol was followed for reduction of the frameworks by *ΔmtrCΔomcA* and *Δbfe.* To assess the amount of Fe(II) contained in biotically-reduced MIL-100, Fe-BTC, and MIL-88A, abiotic and framework-bacteria suspensions were prepared identical to those utilized in reduction kinetics experiments. After 48 h, the framework suspension was centrifuged to pellet the solid, and the supernatant was separated. The isolated solid was washed three times with sterile water and resuspended with SBM using the same volume of initially removed supernatant. Fe(II) concentrations were quantified in the whole sample suspension, isolated supernatant and isolated/washed/resuspended framework. As shown in Figure S7b, the Fe(II) concentrations quantified in isolated supernatant and isolated framework were summed to compare to Fe(II) concentrations measured in the whole sample suspension.

### Cr(VI) adsorption to MR-1 reduced metal-organic frameworks and ferrihydrite

Cr(VI) removal was tested using metal-organic frameworks or ferrihydrite that had been reduced by MR-1 for 24 h. Cultures containing MIL-100 ([Fe(III)]0=15 mM) and 20 mM lactate in SBM were inoculated with MR-1 to OD_600_=0.002, as previously described. After 24 h, K_2_Cr_2_O_7_ was added to each of the cultures ([Cr(VI)]0=70 μM) and Fe(II) concentrations were analyzed. Cr(VI) concentration in the supernatant was analyzed using the DPC assay*(50)*. The DPC assay was performed by mixing 15 μL of sample with 235 μL of DPC solution in a 96-well plate. Absorbance was measured at 540 nm in a microplate reader. DPC solution consisted of DPC (12.5 mg, 103 μmol) in a mixture of 0.5 M H2SO4 (10 mL) and acetone (10 mL). MIL-100 was tested with higher concentrations of Cr(VI) (0.25, 0.50, and 1 mM) using the same procedure.

### Cr(VI) cycling

Cr(VI) removal from a supernatant was tested over ten cycles with MR-1 reduced MIL-100, ferrihydrite, and fumarate. MIL-100, ferrihydrite, and fumarate were biotically reduced for 24 h and K_2_Cr_2_O_7_ was added to each culture ([Cr(VI)]0=0.50 mM). Every 24 h, K_2_cr_2_O_7_ ([Cr(VI)]0=0.50 mM) was added, with ten total additions occurring. Fe(II) concentration and Cr(VI) concentration in the supernatant were monitored as previously described. A second cycling experiment was done using the same conditions, but adding 0.5 mM Cr(VI) until no more Cr(VI) was reduced in the supernatant, equaling 40 additions. To determine cell viability after 10 additions, aliquots of each sample (100 μL) were plated on LB agar plates 24 h after the last Cr(VI) addition and cells were grown aerobically at 30 °C overnight. Following cycling, cycled samples were washed in fresh H_2_O and then dried anaerobically. Prior to STEM imaging, the samples were suspended in ethanol (500 μL) and dropcast on TEM grids. STEM imaging and electron mapping of Fe and Cr was performed using a JOEL 2010F Transmission Electron Microscope.

### Statistical Analysis

Unless otherwise noted, data are reported as mean ± S.D. of N = 3 biological replicates, as this sample size was sufficiently large to detect significant differences in means. Significance was calculated using an unpaired two-tailed Student’s t-test (α=0.05) or ANOVA with post-hoc Dunnett’s test in OriginPro Software (OriginLab, Northhampton, MA).

### H2: Supplementary Materials

Supporting figures and additional methods can be found in the Supplementary Materials.

Materials and Methods

Figure S1. Growth on MIL-88A.
Figure S2. Auto-fluorescence measurements.
Figure S3. Fe(III) reduction in parallel with growth measurements.
Figure S4. Reduction of metal-organic frameworks by MR-1.
Figure S5. Morphology of metal-organic frameworks and ferrihydrite.
Figure S6. Metal-organic framework structural characterization.
Table S1. Metal-organic framework surface areas.
Figure S7. Fe(II) in Biotically Reduced Frameworks.
Figure S8. Reduction of metal-organic frameworks by MtrCAB pathway.
Figure S9. *E. coli* reduction of MIL-100.
Figure S10. Reduction with exogenous riboflavin.
Figure S11. Total Cr removal.
Figure S12. Biotic Cr(VI) reduction.
Figure 13. Cr(VI) cycling and OD_600_ measurements.
Table S2. Cr(VI) cycles.
Table S3. Cr(VI) removal by both biotic and abiotic agents.
Figure S14. Cr(VI) removal for long-term cycling.
Figure S15. Fe(II) reduction of Cr(VI).
Figure S16. Cycled metal-organic framework morphology.
Figure S17. Fe(III) reduction in cycled materials.
Figure S18. Bacterial viability post-cycling.
Figure S19. Stepanovite reduction.

## Acknowledgments

We thank Hazel Mohamedali, Riley Shuping, Gang Fan, and Michael Lucas for their experimental assistance. *S. oneidensis* strains *Δbfe* and *ΔmtrCΔomcA* were generously provided by Prof. Jeffrey Gralnick and *E. coli* MG1655 was generously provided by Prof. Lydia Contreras. We also thank Prof. Contreras for use of her imaging resources. We gratefully acknowledge the use of facilities within the core microscopy lab of the Institute for Cellular and Molecular Biology, University of Texas at Austin and within the X-Ray Diffraction Lab, University of Texas at Austin. S. K. S. thanks the Provost’s Graduate Excellence Fellowship (PGEF) for funding of her PhD scholarship.

## Author Contributions

S.K.S., C.M.D., and B.K.K conceived and designed the experiments. S.K.S. and C.M.D. performed the experiments. S.K.S. performed the material synthesis and characterization. B.K.K. supervised the project. All authors contributed to data analysis and the writing of the manuscript.

## Data availability

All data needed to evaluate the conclusions of this paper are present in the paper and/or Supplementary Materials. Raw data are available from the Texas Data Repository.

