## Supplementary Materials for "Microbial Reduction of Metal-Organic Frameworks Enables Synergistic Chromium Removal"

### Materials and Methods

#### Chemicals and reagents

Trimesic acid ($H_3BTC$, Sigma-Aldrich, 95%), sodium hydroxide ($NaOH$, VWR, pellets), ferrous chloride tetrahydrate ($FeCl_2\cdot 4H_2O$, JT Baker), ferric chloride hexahydrate ($FeCl_3\cdot 6H_2O$, Fisher-Scientific, ACS grade), fumaric acid ($HO_2CCHCHCO_2H$, Alfa Aesar, 99%), red hematite ($Fe_2O_3$, Strem Chemicals, 99.8% Fe), magnesium oxide ($MgO$, Alfa Aesar), oxalic acid ($C_2H_2O_{4-}$, VWR, 10% w/v), HEPES buffer (4-(2-hydroxyethyl)-1-piperazineethanesulfonic acid, IBI Scientific), potassium phosphate dibasic ($K_2HPO_4$, Sigma-Aldrich), potassium phosphate monobasic ($KH_2PO_4$, VWR), sodium chloride ($NaCl$, VWR), ammonium sulfate $(NH_4)_2SO_4$, Fisher Scientific), magnesium(II) sulfate heptahydrate ($MgSO_4\cdot 7H_2O$, Sigma-Aldrich), EDTA acid disodium salt dihydrate ($C_{10}H_{14}N_2Na_2O_8\cdot 2H_2O$, VWR), manganese(II) sulfate monohydrate ($MnSO_4\cdot H_2O$, VWR), ferrous sulfate heptahydrate ($FeSO_4\cdot 7H_2O$, Alfa Aesar), cobalt(II) nitrate hexahydrate ($Co(NO_3)_2\cdot 6H_2O$, Strem Chemicals), calcium chloride dihydrate ($CaCl_2\cdot 2H_2O$, Sigma-Aldrich), zinc(II) sulfate monohydrate ($ZnSO_4\cdot H_2O$, Strem Chemicals), cupric sulfate pentahydrate ($CuSO_4\cdot 5H_2O$, VWR), aluminum potassium sulfate ($AlK(SO_4)_2$, Acros Organics), boric acid ($H_3BO_3$, VWR), sodium molybdate dihydrate ($Na_2MoO_4\cdot 2H_2O$, Beantown Chemical), sodium selenite ($Na_2SeO_3$, Acros Organics), sodium tungstate dihydrate ($Na_2WO_4\cdot 2H_2O$, Alfa Aesar), nickel(II) chloride hexahydrate ($NiCl_{-2}\cdot 6H_2O$, Alfa Aesar), Syto™ 9 green fluorescent nucleic acid stain (ThermoFisher Scientific), SYPRO™ Ruby protein gel stain (ThermoFisher Scientific), and casamino acids (VWR) were used as received. Sodium DL-lactate ($C_3H_5NaO_3$, VWR, 60% in water) was filtered using 0.2 μm PES filters. Sodium fumarate ($Na_2C_4H_2O_4$, VWR) was diluted in $H_2O$ and then filtered using 0.2 μm PES filters. LB Lennox agar powder (VWR) was dissolved in $H_2O$ and sterilized at 121°C and 100 kPa for 1 h. Hydrochloric acid ($HCl$, Sigma-Aldrich, 37%) was diluted in $H_2O$ prior to use. Ferrozine (3-(2-pyridyl)-5,6-bis(4-sulfophenyl)-1,2,4-triazine disodium salt hydrate, TCI Chemicals), ammonium acetate ($NH_4CH_3CO_2$, VWR), hydroxylamine hydrochloride ($HONH_2\cdot HCl$, Alfa Aesar, ACS grade), potassium dichromate ($K_2Cr_2O_7$, Alfa Aesar, ACS grade), 1,5-diphenylcarbazide (DPC, Alfa Aesar), and acetone ($(CH_3)_2CO$, Fisher Scientific, ACS grade) were used as received. Sulfuric acid ($H_2SO_4$, VWR, 95-97%) was diluted in $H_2O$ before use. Iron(III) citrate ($FeC_6H_5O_7$, Alfa Aesar, Fe(III) 16.5-20%, Fe(II) max 5%) was used as received. Deuterium oxide ($D_2O$, Sigma-Aldrich, 99.9%) was used for NMR as received. Nitric acid ($HNO_3$, Sigma-Aldrich, trace mineral grade) was diluted to 2% for ICP-MS analysis. Ultrapure water was produced by a MilliQ Integral Water Purification System.

#### Powder X-Ray Diffraction

PXRD was performed using a Rigaku R-Axis Spider X-Ray Diffractometer with curved image plate detector. The as-synthesized metal-organic frameworks were analyzed without modification. Following reduction of the materials by MR-1, the abiotic and biotic samples were washed with fresh $H_2O$ to remove any residual salts from the medium and stored in anaerobic conditions prior to analysis. Immediately before analysis, the samples were coated in mineral oil to prevent oxidation in the aerobic environment of the instrument. A sample of mineral oil run under the same instrument conditions was subtracted as background from the spectra.

**Surface Area Measurements**

Langmuir surface area measurements were conducted using $N_2$ (99.999%) gas adsorption collected on a Micromeritics ASAP 2020 Physisorption instrument. Approximately 60 mg of either Fe-BTC, MIL-100 or MIL-88A were activated under high vacuum at 120 °C with a ramp of 0.1 deg/min for 24 h before analysis.

**Electron Microscopy**

SEM was used to determine the extent of aggregation in the as-synthesized metal-organic framework. Images were collected using Zeiss Supra 40V SEM. TEM was used to determine particle morphology of cycled MIL-100. The TEM grids used for STEM and element mapping were imaged using a Thermo Fisher Tecnai TEM.

**Leaching and Framework stability analysis**

The metal-organic frameworks were tested for leaching and framework stability by exposing the materials to culture conditions for 48 h. The ferrozine assay was used to determine the Fe(III) concentration in the supernatants after the metal-organic frameworks ([Fe(III)=15mM). For Fe(II) analysis, samples were prepared as previously described. For total Fe analysis, the sample was mixed with 1 M hydroxylamine hydrochloride in a 1:1 ratio and analyzed by the ferrozine assay. Fe(III) concentrations were calculated by subtracting the Fe(II) concentration from the total Fe concentration.

NMR spectroscopy was used to determine the extent of fumarate leaching from MIL-88A ([Fumarate$^-$]=5 mM) in 1 mL of $D_2O$ and stored anaerobically at 30 °C. After 48 h, the samples were centrifuged and the supernatant was removed. The supernatant was spiked with 1.0 mg of benzoic acid as an internal standard. A standard of fumaric acid (1.0 mg/mL) and benzoic acid (1.0 mg/mL) was mixed in 1 mL $D_2O$. An aliquot of the supernatant (50 µL) was diluted in 950 µL of $D_2O$ and analyzed on an Agilent MR400 NMR (400 MHz).

Framework structural stability was tested by exposing the metal-organic frameworks to culture conditions for 48 h. Following exposure, the materials were washed and analyzed by PXRD.

**Auto-fluorescence measurements**

To determine the viability of an auto-fluorescence assay in anaerobic conditions, cultures containing fumarate (40 mM) and lactate (20 mM) in SBM were inoculated with stationary phase MR-1 ($OD_{600}$=0.002, anaerobic pregrowth) and incubated at 30 °C in anaerobic conditions. For analysis, an aliquot of the culture (250 μL) was transferred to a 96-well plate. Fluorescence was measured anaerobically. Background was subtracted from all fluorescence measurements and experiments were done in triplicate. To confirm that auto-fluorescence trends matched those of optical density, $OD_{600}$ was measured simultaneously by loading 2 μL of sample onto a ThermoFisher NanoDrop 2000c and measuring absorbance at 600 nm. Background was subtracted from all absorbance measurements. Absorbance measurements were corrected by a factor of 10 to account for using the microvolume sampler.

Auto-fluorescence of MR-1 grown on ferrihydrite was also measured. Cultures containing ferrihydrite (15 mM) and lactate (20 mM) in SBM were inoculated with stationary phase MR-1 ($OD_{600}$=0.002, anaerobic pregrowth). At each time point, an aliquot of the sample was removed and gently spun down for 1 minute to pellet the ferrihydrite. The supernatant (250 μL) was transferred to a 96-well plate and fluorescence was assessed as previously described. All experiments were done in triplicate.

Because metal-organic frameworks are known for adsorption properties, we ensured that the materials were not interacting with the secreted flavins or quenching their auto-fluorescence. To test this, each metal-organic framework ([Fe(III)=15 mM) was mixed with 20 mM lactate and 10 μM riboflavin in SBM. A control sample of 20 mM lactate and 10 μM riboflavin in SBM was also tested. Samples were stored in the dark in anaerobic conditions at 30 °C for 24 h. To measure fluorescence, an aliquot of each sample was gently centrifuged and the supernatant (250 μL) was transferred to a 96-well plate. Fluorescence was measured as previously described. All experiments were done in triplicate.

**Reduction of MIL-100 by *E. coli***

To demonstrate that the reduction of MIL-100 is not due to promiscuous reduction, a culture of MIL-100 ([Fe(III)]=15 mM) and 20 mM lactate in SBM was inoculated with stationary-phase *E. coli* ($OD_{600}$=0.002). *E. coli* MG1655, generously provided by Dr. Lydia Contreras (University of Texas, Austin, TX), was pregrown anaerobically in 40 mM fumarate and 20 mM lactate in SBM, washed twice and diluted to an $OD_{600}$=0.2 prior to culture inoculation. All cultures were incubated at 37 °C in an anaerobic environment.

Fe(II) concentrations were analyzed using the ferrozine assay. An abiotic control was also tested. Experiments were performed in triplicate.

#### Fe(II) concentrations of MR-1 reduced MIL-100 supernatant

Total Fe(II) concentration and Fe(II) concentration in the supernatant was determined for MR-1 reduced MIL-100 cultures. Cultures containing MIL-100 ([Fe(III)]=15 mM) and 20 mM lactate in SBM were inoculated with stationary phase MR-1 ($OD_{600}$=0.002; anaerobic pregrowth). For total Fe(II) analysis, the cultures were gently mixed to suspend the MIL-100 particles and a 20 µL aliquot was removed. For supernatant Fe(II) analysis, an aliquot of the culture was gently spun down to pellet the material and 20 µL of the supernatant was removed. The 20 µL aliquots were analyzed using the ferrozine assay. An abiotic control was also tested.

#### ICP-MS analysis

ICP-MS was used to analyze total Cr in the supernatant of biotically-reduced MIL-100 challenged with 0.5 mM Cr(VI). Cultures containing MIL-100 ([Fe(III)=15 mM) and 20 mM lactate were inoculated with MR-1 ($OD_{600}$=0.002, anaerobic pregrowth). MR-1 reduced MIL-100 until the [Fe(II)] = 2.8 mM after which cultures were challenged with 0.5 mM Cr(VI) and aliquots of the supernatant were removed for analysis. Samples were centrifuged (10000 x g, 1 min), frozen in liquid $N_2$ and stored at -20°C until further sample preparation. For the ICP-MS sample preparation, the samples were thawed and diluted 200-fold in 2% $HNO_3$ immediately prior to analysis. A 7500ce Agilent ICP-MS was used to analyze total Cr (LOD:0.54 ppb) and was verified using m/Z=52 and 53.

#### MR-1 reduction of Cr(VI)

Reduction of 100 µM Cr(VI) by MR-1 was tested by inoculating a culture of 20 mM lactate and 50 µM $K_2Cr_2O_7$ in SBM with stationary phase MR-1 ($OD_{600}$=0.002, anaerobic pregrowth). Cultures were stored at 30 °C in anaerobic conditions for 24 h. Cr(VI) concentrations were analyzed using the DPC assay. An abiotic control was also tested.

#### Cr(VI) reduction by $FeCl_2$

Cr(VI) reduction by abiotic Fe(II) was tested by mixing a solution of $FeCl_2 \cdot 4 \cdot H_2O$ (either 0.5 mM, 1.0 mM or 2.0 mM), 20 mM lactate and 0.5 mM Cr(VI) in SBM (pH=7.2). Solutions were stored in anaerobic conditions at 30 °C for 24 h. Cr(VI) concentrations were determined using the DPC assay.

#### Cr(VI) reduction by MR-1 reduced Fe(III)-citrate

Cr(VI) reduction by biotically reduced Fe(II) was tested by inoculating a culture of Fe(III)-citrate (5 mM) and 20 mM lactate in SBM with stationary phase MR-1 ($OD_{600}$=0.002, anaerobic pregrowth). MR-1 reduced Fe(III)-citrate until Fe(II) concentrations for 8 h, or when Fe(II) concentrations were comparable to those observed at 24 h with MR-1 and MIL-100. Fe(II) concentration in the Fe(III)-citrate and MR-1 cultures was 2.5 mM after 8 h. Once Fe(III) reduction was established, the cultures were challenged with 0.5 mM Cr(VI). After 12 h, the cultures were challenged again with 0.5 mM Cr(VI). Fe(II) and Cr(VI) concentrations were monitored throughout the experiment using the ferrozine and DPC assays, respectively. An abiotic control was also tested using the same procedure.

**MR-1 reduction of stepanovite**

Fe(III) reduction in a natural metal-organic framework, stepanovite, was tested by inoculating a culture of stepanovite ($[Fe(III)]_0$=15 mM) and 20 mM lactate in SBM with stationary phase MR-1 ($OD_{600}$=0.002, anaerobic pregrowth). Fe(II) concentrations were analyzed using the ferrozine assay. As stepanovite contains a carbon source (oxalate) for MR-1, Fe(III) reduction in a culture containing just stepanovite ([Fe(III)]=15 mM) in SBM was inoculated with stationary phase MR-1 ($OD_{600}$=0.002, anaerobic pregrowth). Fe(II) concentrations were analyzed using the ferrozine assay. Abiotic controls were also tested.

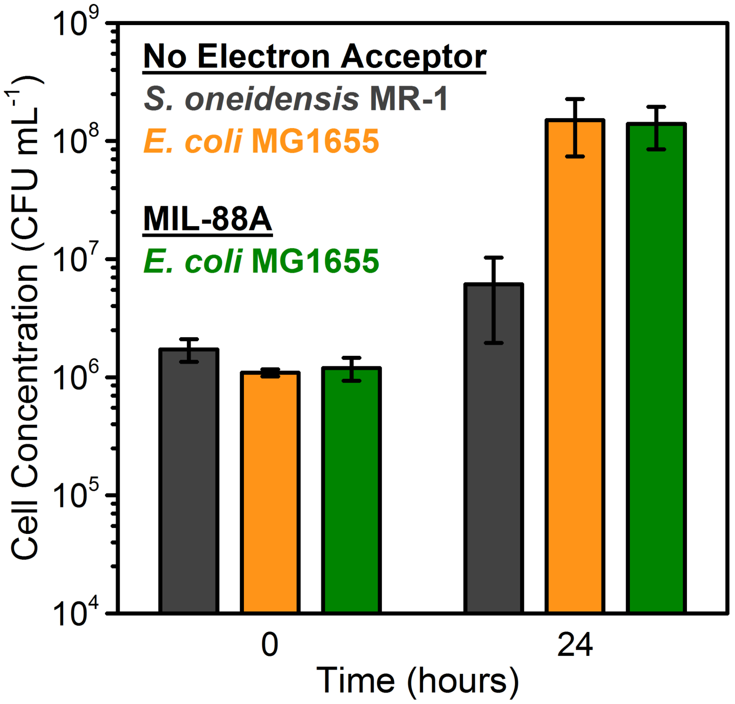


**Figure S1. Growth on MIL-88A.** CFU counts of *S. oneidensis* MR-1 and *E. coli* MG1655 with no electron acceptor present (only SBM and lactate), as well as MG1655 with MIL-88A as an electron acceptor. The initial inoculating $OD_{600}$=0.002. Data show mean ± S.E. for three independent biological replicates.

**a**

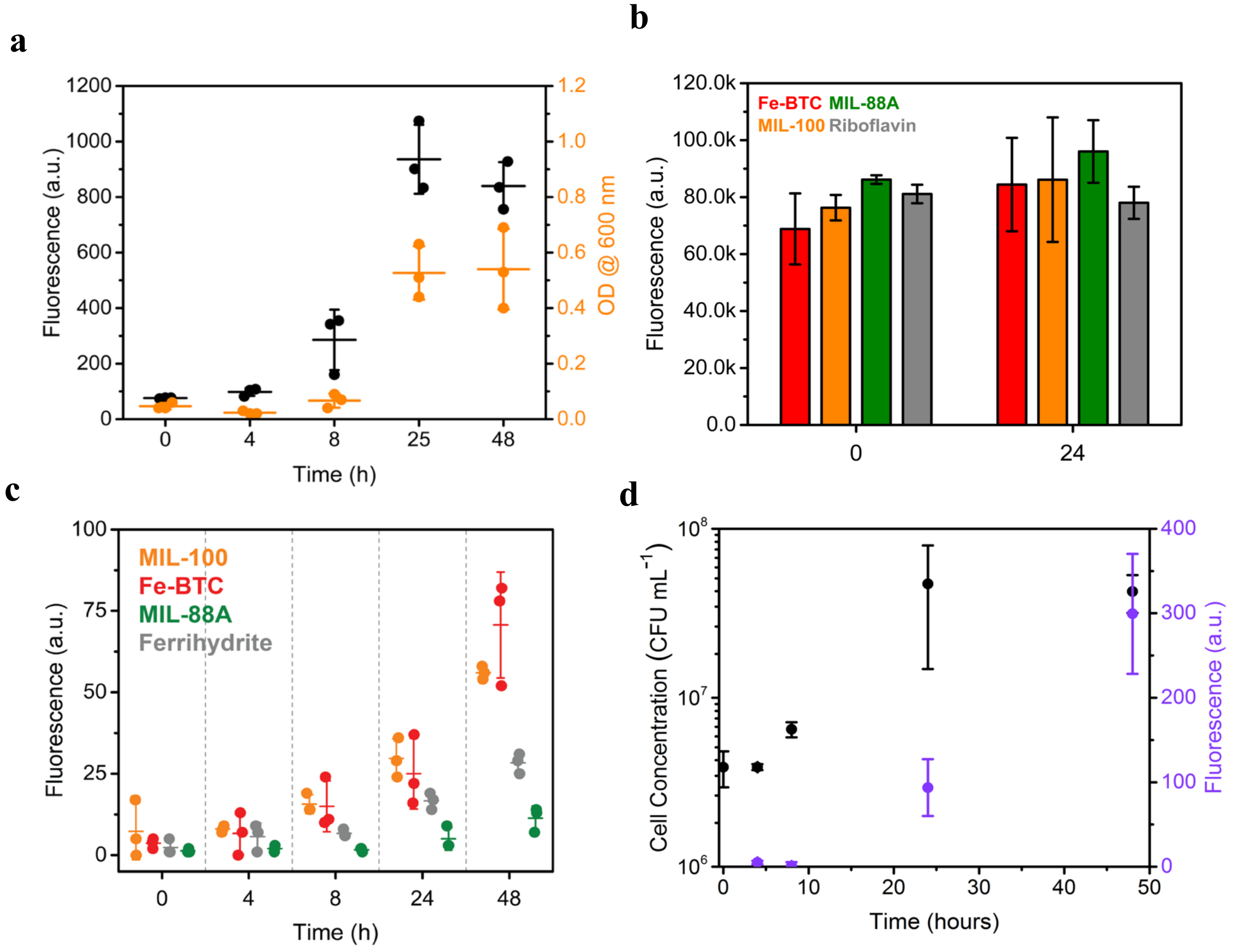


**Figure S2. Auto-fluorescence measurements.** (a) Auto-fluorescence (*black*) and $OD_{600}$ (*orange*) of MR-1 (inoculating $OD_{600}$=0.002) cultured with 40 mM fumarate and 20 mM lactate in SBM. Data show mean ± S.D. for three independent biological replicates. (b) Fluorescence of 10 µM riboflavin when mixed with Fe-BTC, MIL-100, MIL-88A, and no material at 0 h and 24 h. Data show mean ± S.D. for three independent replicates. (c) Background subtracted auto-fluorescence of MR-1 when cultured with MIL-100, Fe-BTC, MIL-88A, and ferrihydrite over 48 h. For both Fe-BTC and MIL-100, there is a significant difference between the auto-fluorescence of 24 h and 48 h ($P < 0.01$). Data show mean ± S.D. for three biological replicates. (d) CFU counts (black) and auto-fluorescence (purple) of MR-1 grown on MIL-100 over 48 h. Data show mean ± S.E. for three independent biological replicates.

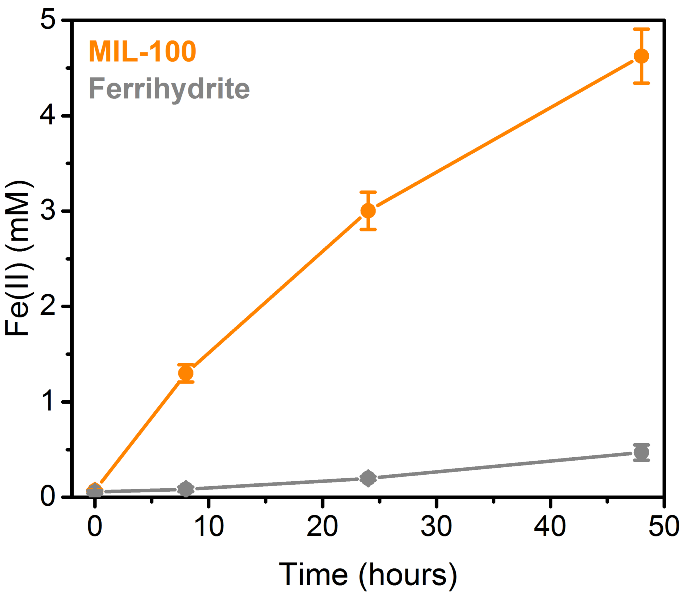


**Figure S3. Fe(III) reduction in parallel with growth measurements.** Fe(III) reduction by MR-1 grown on MIL-100 and ferrihydrite over 48 h, measured in parallel with Syto™ 9 and SYPRO™ Ruby (Figure 2d-e). The initial inoculating $OD_{600}$=0.02. Data show mean ± S.D. for three independent biological replicates.

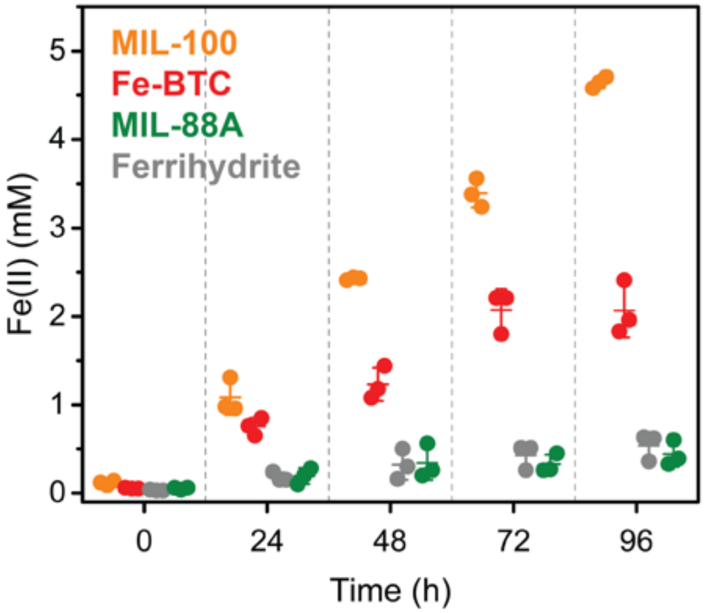


**Figure S4. Reduction of metal-organic frameworks by MR-1.** Reduction of MIL-100, Fe-BTC, MIL-88A, and ferrihydrite by MR-1 over 96 h. The initial inoculating $OD_{600}$=0.002. Data show mean ± S.D. for three independent biological replicates.

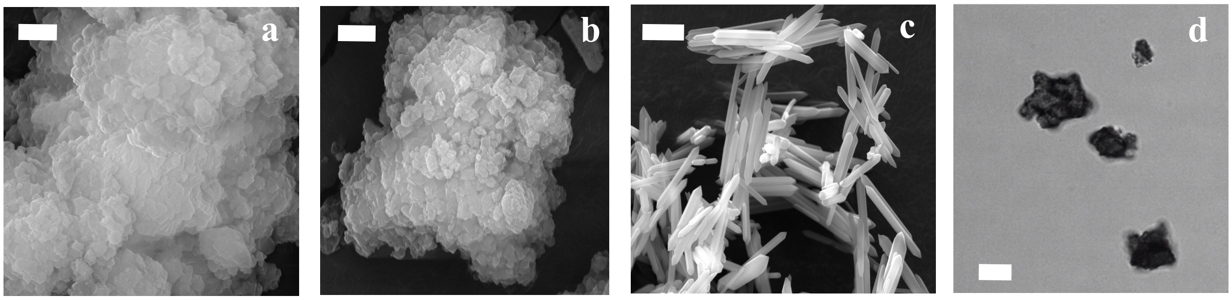


**Figure S5. Morphology of metal-organic frameworks and ferrihydrite.** SEM images of (a) MIL-100, (b) Fe-BTC, and (c) MIL-88A as synthesized. Scale bars are 2 µm. (d) Brightfield microscopy of ferrihydrite. Scale bar is 10 µm.

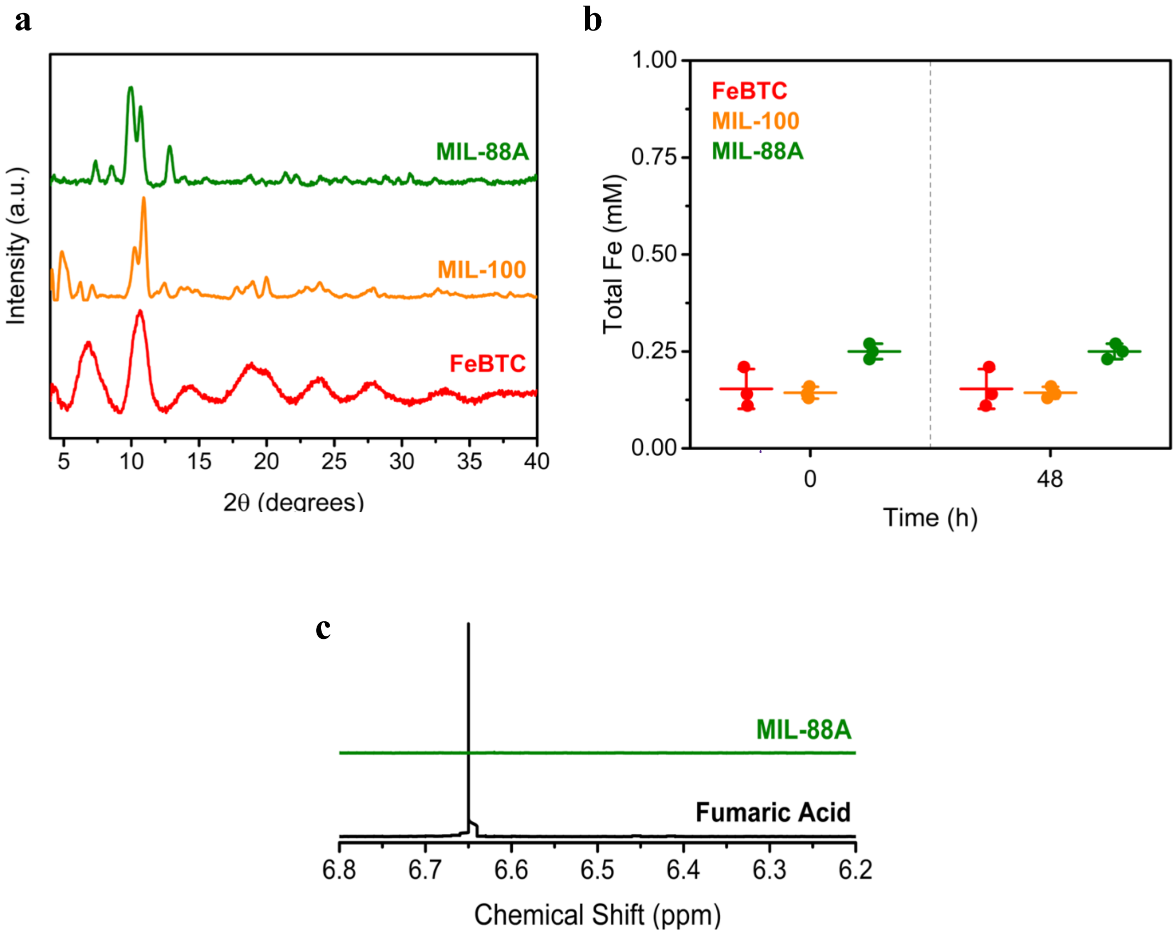


**Figure S6. Metal-organic framework structural characterization.** (a) PXRD spectra of Fe-BTC, MIL-100, and MIL-88A after exposure to standard culture conditions for 48 h (SBM, 20 mM lactate 30 ºC). (b) [Fe(III)] in the supernatant of Fe-BTC, MIL-100, and MIL-88A after 48 h. (c) $^{1}H$ NMR of the MIL-88A supernatant after 48 h and a 1 mg/mL fumaric acid standard.

**Table S1. Metal-organic framework surface areas.**

| Material | Measured Langmuir Surface Area ($m^2g^{-1}$) | Reported BET Surface Area ($m^2g^{-1}$) | Reference |
|---|---|---|---|
| **MIL-100** | 2188 ± 25 | 1974 | (*45*) |
| **Fe-BTC** | 1512 ± 92 | 1092 | (*46*) |
| **MIL-88A** | 130 ± 3 | 25 | (*47*) |

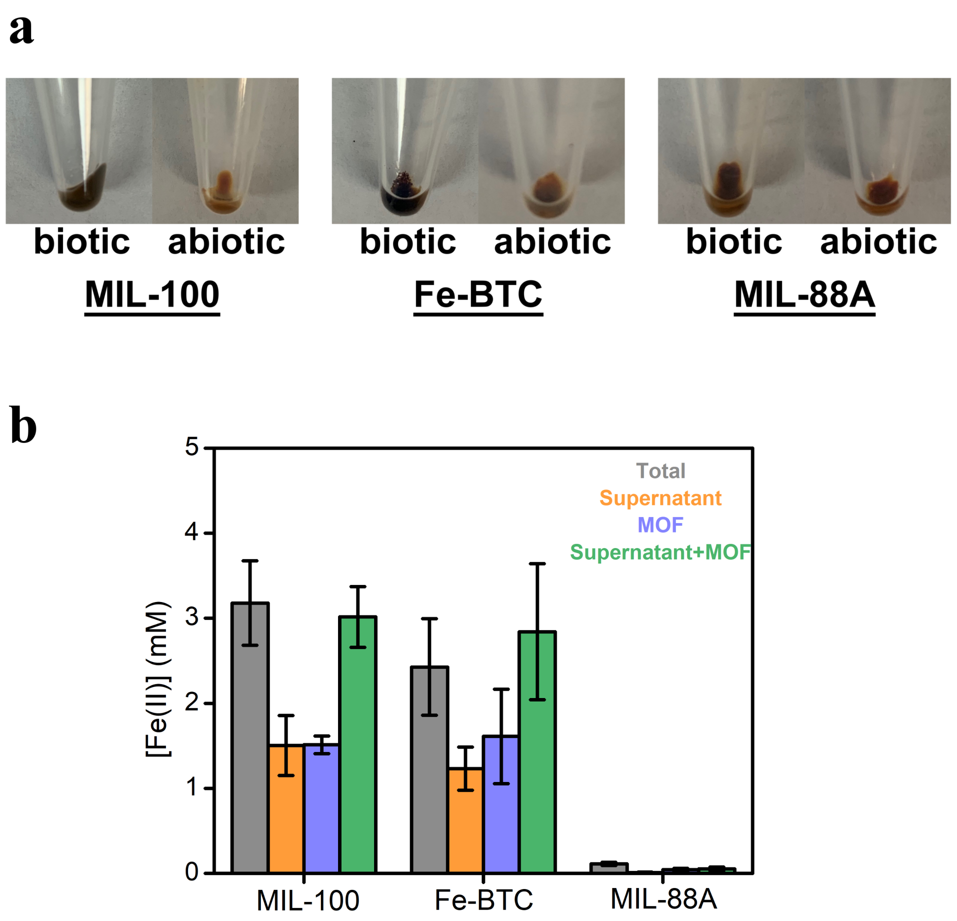


**Figure S7. Fe(II) in Biotically Reduced Frameworks.** (a) Images of biotic and abiotic treated MIL-100, Fe-BTC, and MIL-88A after 48 h. (b) Fe(II) concentrations as measured in the isolated supernatant (*orange*), isolated framework (*blue*), and whole sample suspension (*grey*). Green bars show the addition of the measured isolated supernatant and isolated framework concentrations. Data show mean ± S.D. for three independent experiments.

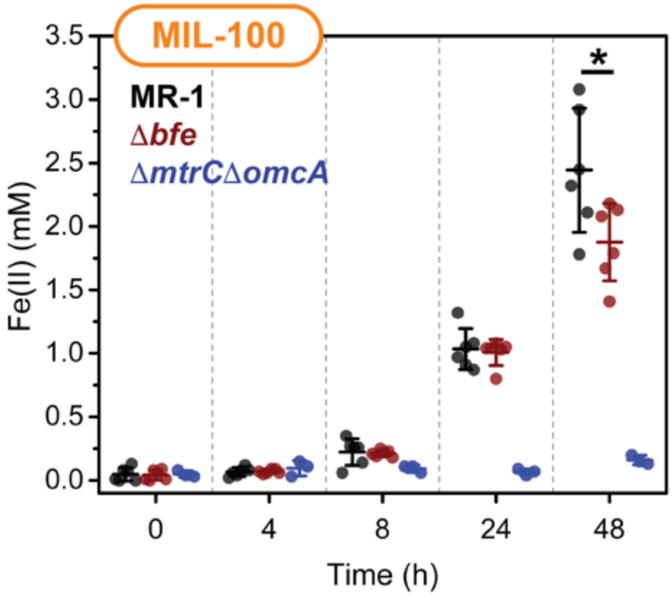


**Figure S8. Reduction of metal-organic frameworks by MtrCAB pathway.** Reduction of (a) MIL-100 by MR-1, Δ*bfe,* and Δ*mtrC*Δ*omcA* over 48 h. Data show mean ± S.D. for six independent experiments. *P<0.05, unpaired two-tailed *t*-test.

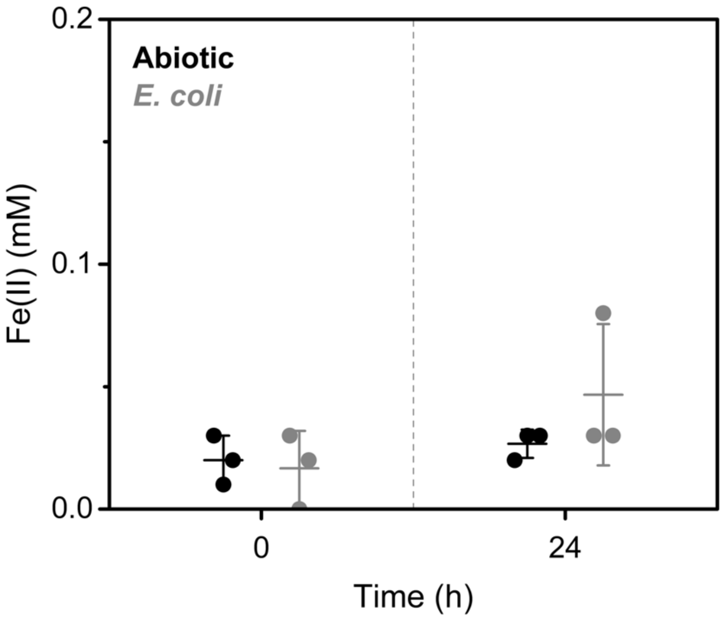


**Figure S9. *E. coli* reduction of MIL-100.** Reduction of Fe(III) by *E. coli* (inoculating $OD_{600}$=0.002) when cultured with MIL-100 ($[Fe(III)]_0$=15 mM) and 20 mM lactate in SBM.

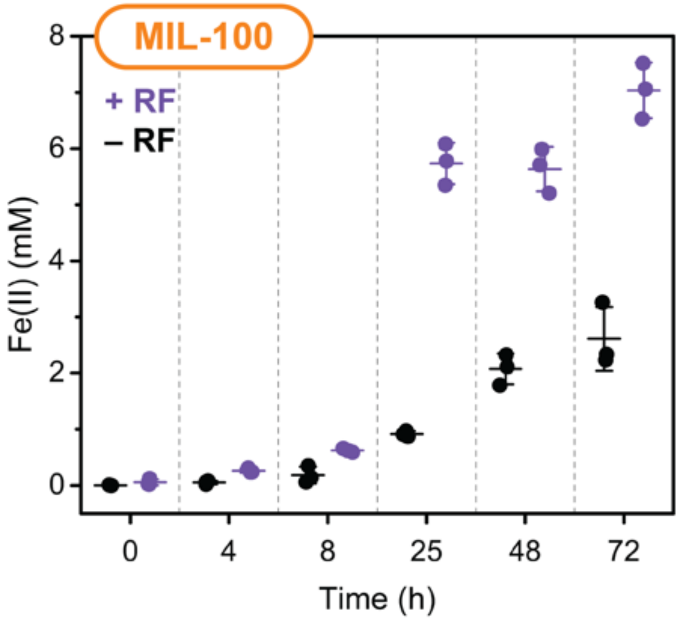


**Figure S10. Reduction with exogenous riboflavin.** Reduction of MIL-100 by MR-1 with and without exogenous 10 µM riboflavin supplementation. Data show mean ± S.D. for three independent experiments.

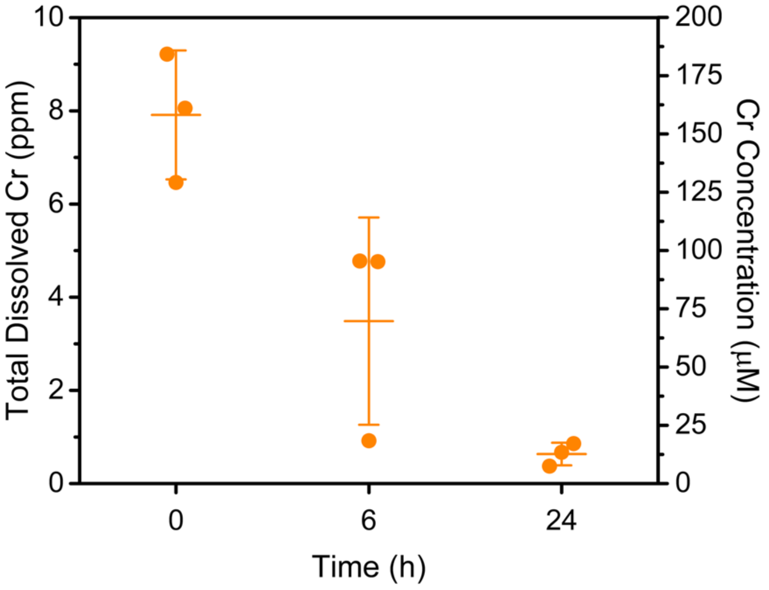


**Figure S11. Total Cr removal.** Total Cr, as measured by ICP-MS, in the supernatant of MR-1 reduced MIL-100 after a challenge with 0.5 mM Cr(VI).

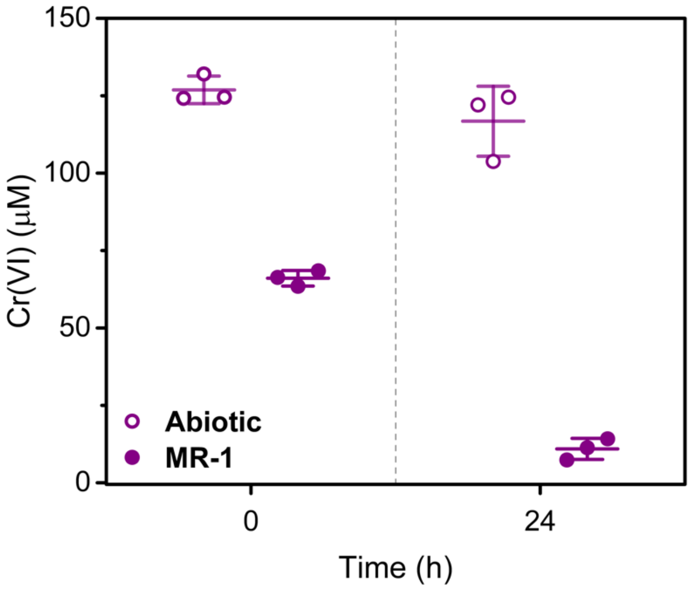


**Figure S12. Biotic Cr(VI) reduction.** Reduction of Cr(VI) by MR-1 in the presence of 100 μM Cr(VI)

a

b

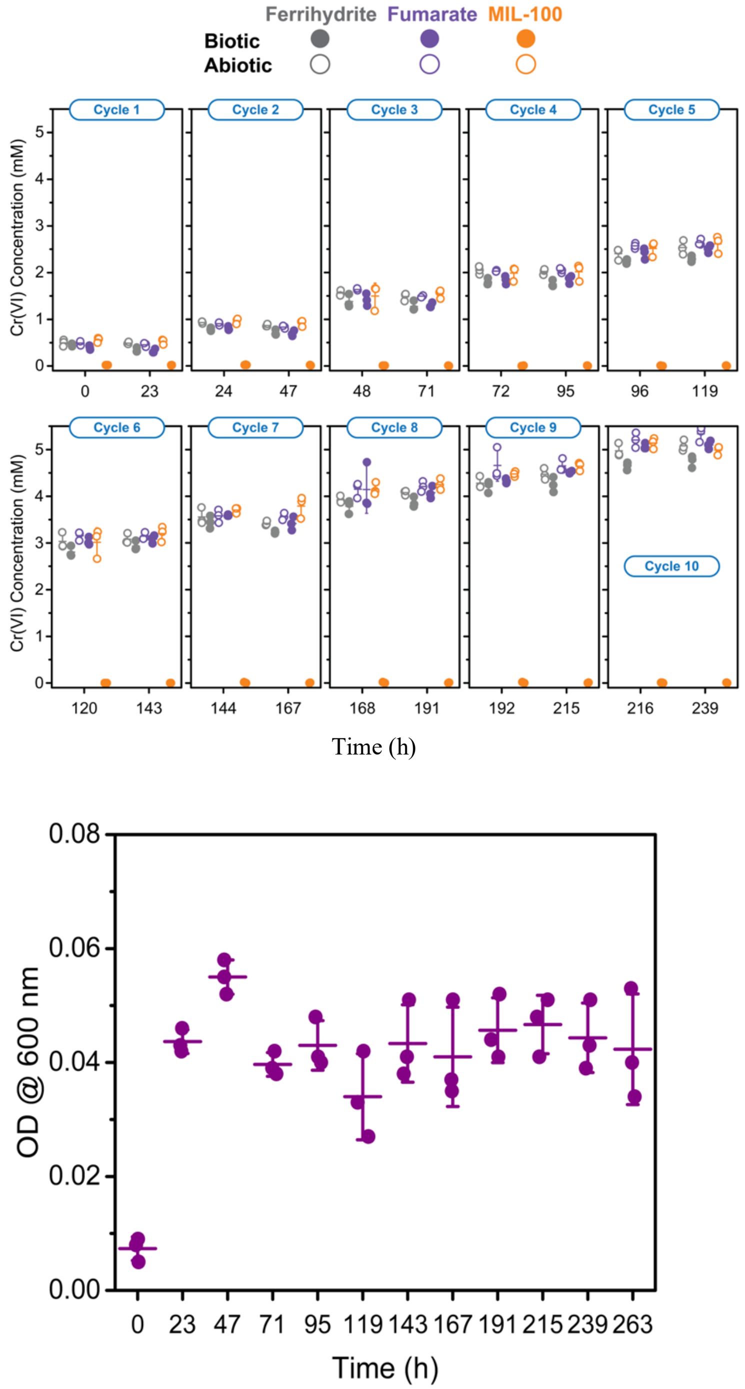


**Figure S13. Cr(VI) cycling and $OD_{600}$ measurements.** (a) Cr(VI) concentrations in the supernatant for each Cr(VI) challenge (1-10) for MIL-100 (*orange*), ferrihydrite (*grey*), and fumarate (*purple*). (b) Growth of MR-1 when grown on soluble fumarate (40 mM) over the course of the 10 Cr(VI) cycles.

**Table S2. Cr(VI) cycles.**

| Cycle Number | Culture Volume (μL) | Volume of 500 mM Cr(VI) Added (μL) | Cr(VI) Removed (mg $g^{-1}$) Biotic MIL-100 | Cr(VI) Removed (mg $g^{-1}$) Biotic Ferrihydrite |
|---|---|---|---|---|
| **1** | 1960 | 1.96 | 7.61 ± 0.09 | 3.28 ± 1.07 |
| **2** | 1842 | 1.84 | 15.49 ± 0.09 | 5.96 ± 1.25 |
| **3** | 1724 | 1.72 | 23.52 ± 0.09 | 4.59 ± 2.20 |
| **4** | 1606 | 1.61 | 31.40 ± 0.09 | 5.00 ± 1.48 |
| **5** | 1487 | 1.49 | 39.23 ± 0.09 | 4.25 ± 1.34 |
| **6** | 1369 | 1.37 | 47.26 ± 0.00 | 1.37 ± 2.08 |
| **7** | 1250 | 1.25 | 55.09 ± 0.09 | 5.55 ± 0.62 |
| **8** | 1131 | 1.13 | 62.97 ± 0.09 | 2.74 ± 2.25 |
| **9** | 1012 | 1.01 | 70.85 ± 0.09 | 5.21 ± 3.29 |
| **10** | 893 | 0.89 | 78.67 ± 0.09 | 5.14 ± 2.56 |

**Table S3. Cr(VI) removal by both biotic and abiotic agents.**

| Material | Cr(VI) Capacity (mg $g^{-1}$) | Process | Reference |
|---|---|---|---|
| **Humic Acid Coated Magnetite Nanoparticles** | 3.37 | Fe Oxidation | (*51*) |
| **Bio amended $Fe^{0}$** | 6.2 | Fe Oxidation | (*52*) |
| **Ferrihydrite** | 12.97 | Fe Oxidation | (*53*) |
| **FIR-53** | 17.8 | Anion Exchange | (*34*) |
| ***Geobacter sulfurreducens*-Reduced Chlorite (+AQDS)** | 20.8 | Fe Oxidation | (*31*) |
| **Sulfamate-Bacterial Cellulose** | 22.73 | $-NH_3$ and $–OH_2$ Reactions | (*54*) |
| **Living *Bacillus coagulans* Biomass** | 23.8 | Biosorption | (*55*) |
| **Wheat Straw Biochar** | 24.6 | Surface Reduction | (*56*) |
| **FIR-54** | 24.7 | Anion Exchange | (*34*) |
| **Aluminum Substituted Ferrihydrite** | 39.79 | Fe Oxidation | (*54*) |
| **Dead *B. coagulans* Biomass** | 39.9 | Biosorption | (*55*) |
| **ABT·$2ClO_4$** | 65 | Anion Exchange | (*32*) |
| **MONT-1** | 55.2 | Anion Exchange | (*32*) |
| **Zn-MOF-74** | 750 | Fe(II) Surface Enhancement | (*35*) |
| **UTSA-74** | 796 | Fe(II) Surface Enhancement | (*35*) |

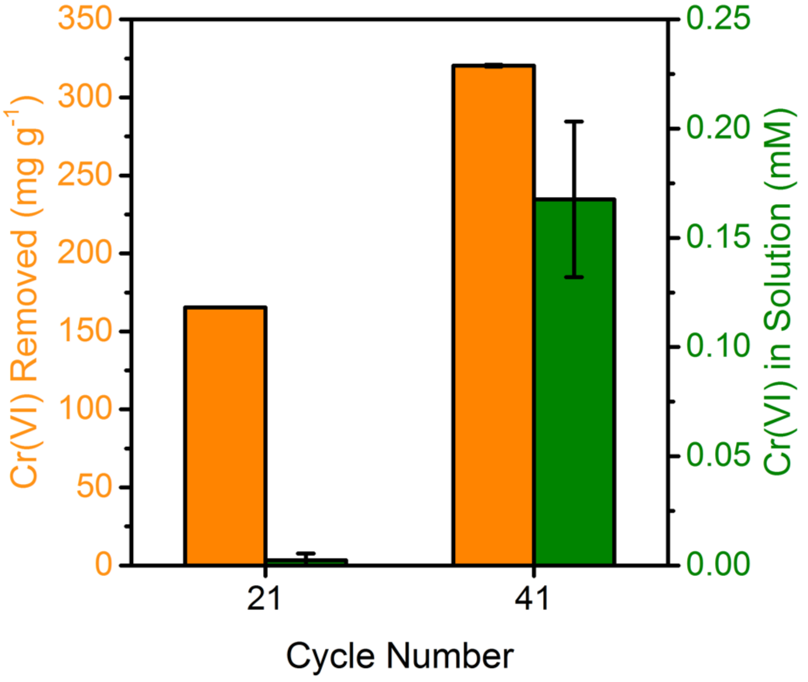


**Figure S14. Cr(VI) removal for long-term cycling.** Cr(VI) remaining in the supernatant and total Cr(VI) removed for biotic MIL-100 samples after 21 and 41 additions of 0.5 mM Cr(VI).

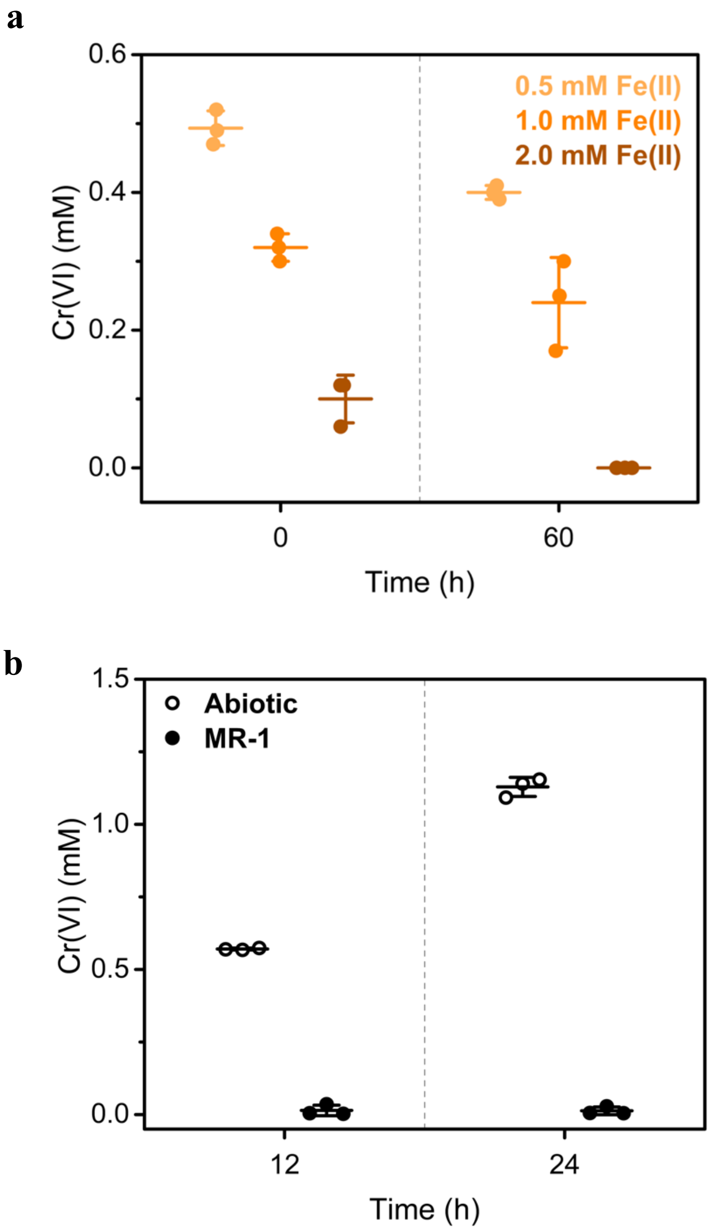


**Figure S15. Fe(II) reduction of Cr(VI).** Reduction of 0.5 mM Cr(VI) by (a) $FeCl_2 \cdot 4H_2O$ (either 0.5 mM, 1.0 mM, or 2.0 mM) after 24 h and by (b) MR-1 reduced Fe(III)-citrate (2 cycles). MR-1 was grown on Fe(III)-citrate for ca. 12 h prior to initial Cr(VI) addition. A second dose of Cr(VI) was added at 24 h.

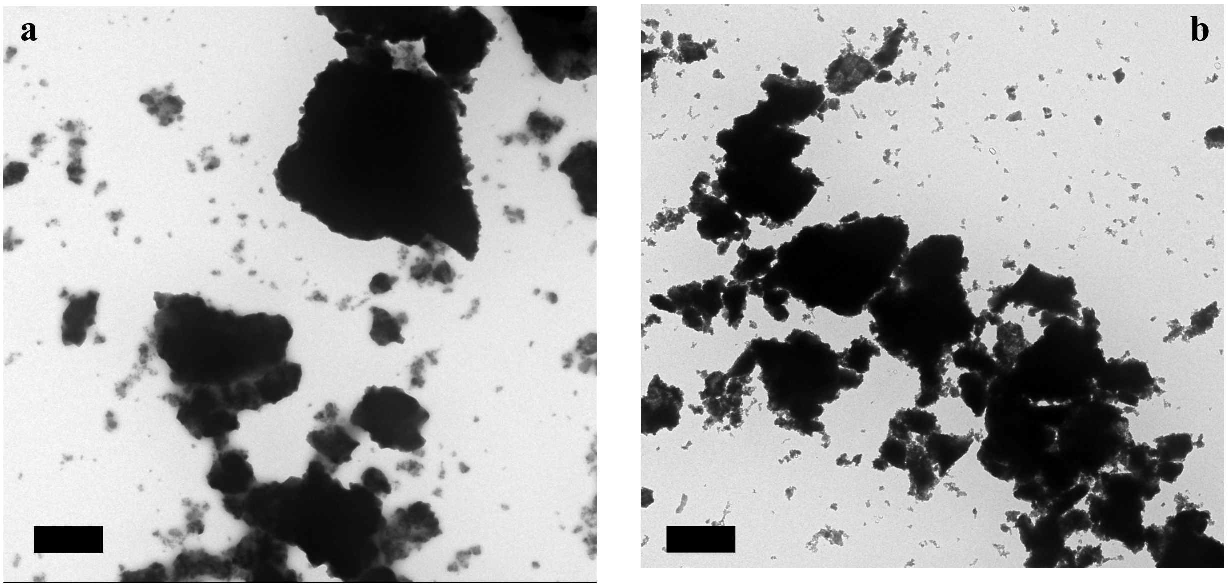


**Figure S16. Cycled metal-organic framework morphology.** TEM images of (a) abiotic MIL-100 and (b) biotic MIL-100 after 10 Cr(VI) challenges. Scale bars are 1 μm.

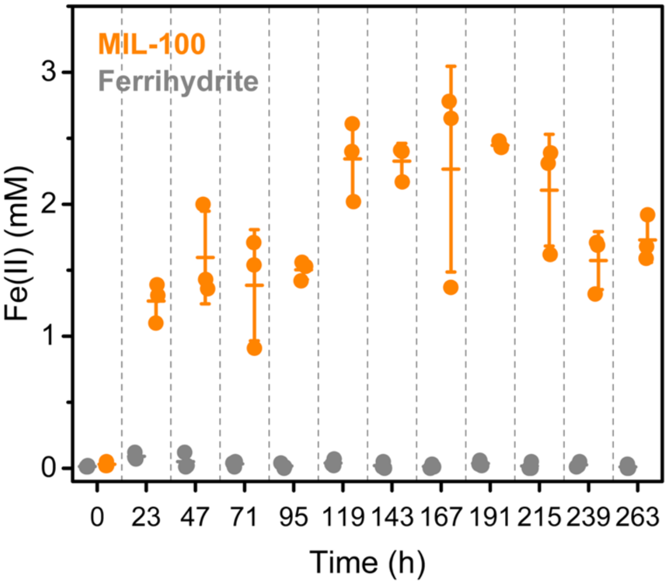


**Figure S17. Fe(III) reduction in cycled materials.** Fe(III) reduction of ferrihydrite (*grey*) and MIL-100 (*orange*) over the course of 10 Cr(VI) challenges.

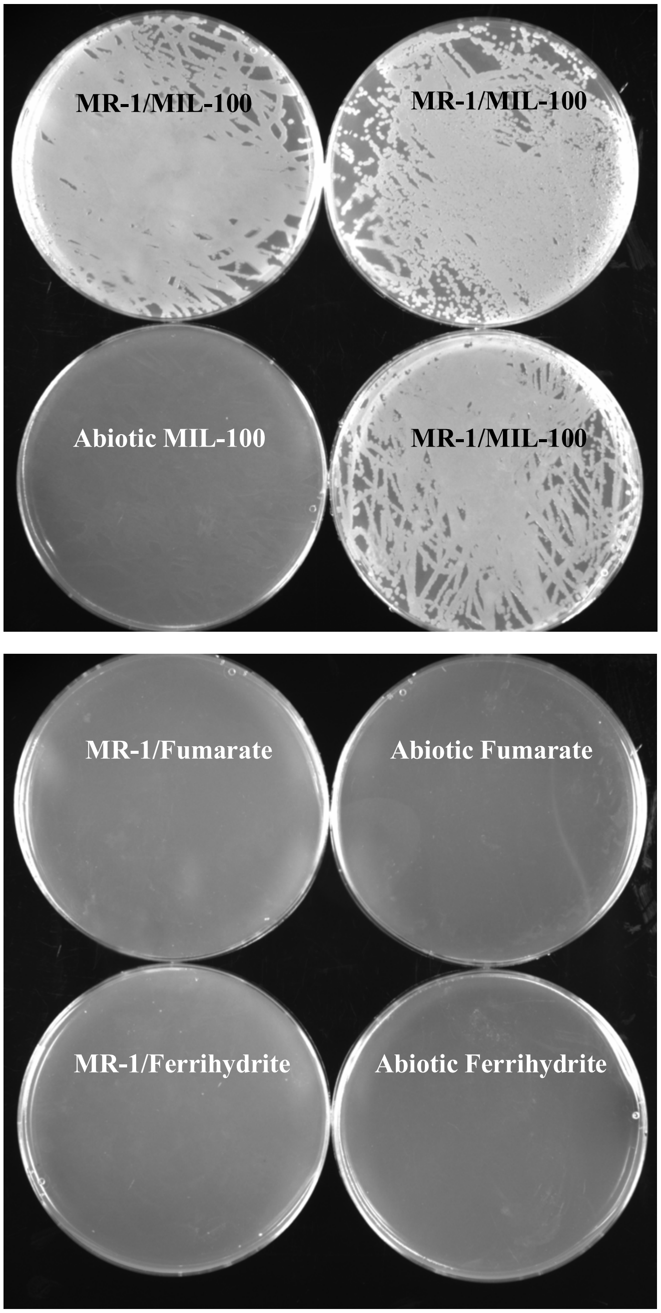


**Figure S18. Bacterial viability post-cycling.** Bacterial viability assessed by plating on LB agar plates 23 h after the tenth 0.5 mM Cr(VI) challenge in MR-1/MIL-100, MR-1/Ferrihydrite, MR-1/Fumarate, and the abiotic samples. MIL-100 biotic samples are shown in triplicate, while only a single plate is shown for the others. Only MR-1/ MIL-100 samples exhibited growth.

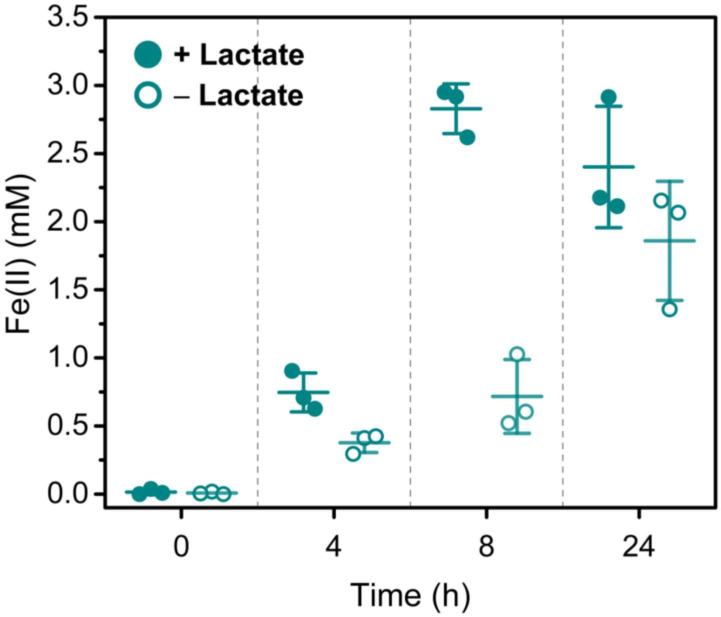


**Figure S19. Stepanovite reduction.** Reduction of stepanovite with and without an added carbon source (lactate) by MR-1.
